# Elimination of intramuscular immunoglobin accumulation alleviates Duchenne Muscular Dystrophy

**DOI:** 10.64898/2026.02.11.703666

**Authors:** Lei Zhang, Wenting Guo, Mei Huang, Zhixin Liang, Enni Zhu, Geng Li, Chong Chang, Xuan Guo, Wenjie Sun, Jia Li, Jie Liu, Han Qiu, Lizhen Zhu, Shuaiwei Ren, Raoxian Bai, Jiaojian Wang, Yinping Gao, Yan Song, Fubin Li, Yi Dai, Zhiying Wu, Jing Hu, Weizhi Ji, Yongchang Chen, Xing Chang

## Abstract

Duchenne muscular dystrophy (DMD) is a devastating neuromuscular disorder due to loss of dystrophin, a cytoskeletal protein critical for muscle integrity and functionality. Despite recent therapeutic advances, there remains a significant unmet need for more effective and accessible therapeutics. Here, we discovered an early accumulation of immunoglobulin G (IgG) in the sarcolemma, which exacerbates tissue inflammation and disease progression. The IgG accumulation primarily resulted from ectopic localization of FcγR1, a high-affinity IgG Fc receptor, on dystrophin-deficient myofibers. In two independent murine models, eliminating IgG accumulation via B cell depletion provided sustained benefit to alleviate DMD progression. Our findings uncover a novel disease accelerator in DMD and demonstrate the potential to target this mechanism as therapeutics for broader population of patients with DMD.

## Introduction

Duchenne muscular dystrophy (DMD) is the most common childhood form of muscular dystrophy, affecting one in 3500 to 5000 males at birth globally (*1, 2*). Patients with DMD suffer a progressive wasting of both skeletal and cardiac muscles due to the loss of musculoskeletal protein dystrophin encoded by the *DMD* gene on chromosome Xp21(*3, 4*). Despite extensive research, this disease remains incurable as patients often decease from progressive muscle damage and cardiorespiratory failure in early thirties(*5*). While substantial therapeutic development have been made to restore truncated dystrophin via exon skipping (*6–12*) or gene therapies (*13–15*), these approaches only benefit a limited patient sub-population with defined genetic alterations. In contrast, targeting key common pathological event(s) in DMD holds the potential to benefit a broader range of patients with DMD, regardless of their specific mutation(s) in DMD gene or progression stages (*16*).

Although inflammation is known to be associated with DMD, effective and safe therapies to curb the inflammatory responses remain to be developed. This is partially due to the multifaceted complexity of the immune system and incomplete understanding of the driving immune mechanisms in DMD (*1, 17*). Current research suggests that the absence of dystrophin compromises the membrane integrity of muscle fibers, rendering them susceptible to contraction-induced injury. This vulnerability ultimately leads to inflammatory cell death (*e.g.* necrosis) together with a profound inflammatory response (*18*). However, previous efforts targeting inflammatory effectors using ciclosporin A(*19*), anti-TNFα(*20*), IL-1β antagonist (*21*), Edasalonexent (a NF-κb inhibitor)(*22*), and Flavocoxid (*23*) only resulted in insignificant clinical benefits in patients with DMD (*24*), casting doubt on the importance of immune system in the pathogenesis of DMD.

In this study, we demonstrate that ectopic accumulation of immunoglobin at the plasma membrane of myofibers plays a critical role in DMD pathogenesis. This accumulation was primarily caused by upregulation of FcγR1 (CD64), a high-affinity receptor for IgG in dystrophin-deficient myofibers. Importantly, reducing antibody accumulation - achieved through anti-CD20-mediated B cell depletion - promoted a transition from a pro-inflammatory environment to a pro-healing state in DMD muscles. Consequently, anti-CD20 antibody treatment alleviated muscular dystrophy and provide long-term benefits to improve skeletal muscle functions in two murine models of DMD. Our findings reveal a previously unknown ectopic immunoglobin accumulation and its downstream molecule events that accelerate DMD progression. We further highlight the potential of targeting B cells and/or immunoglobin accumulation as a therapy for this currently incurable disease.

## Results

### Accumulation of immunoglobulins at the sarcolemma of skeletal muscles in DMD animal models

In the previous characterization of the *Dmd*^E4*^ mice, which contains a 4-bp deletion in exon 4 of the *Dmd* gene(*12*), ectopic accumulation of immunoglobin G (IgG) was observed in various muscle tissues. As early as 2-weeks after birth, IgG was detected in the skeletal muscles of *Dmd*^E4*^ mice, including tibialis anterior (T.A.) (**Figure 1A, fig. S1A**), rectus femoris (R.F.) (**fig. S1B**), diaphragm (**fig. S1C**), but not in cardiac muscles (**Fig. S1D**). IgG was primarily deposited at the sarcolemma, evidenced by its co-localization with laminin (**Figure 1A**), a cell-adhesion molecule localized to the basement membrane of skeletal muscles. In contrast to IgG, the other abundant antibody isotype in the serum, IgM was absent in the muscles of 2-week-old *Dmd*^E4*^ mice (**fig. S1, A to D**). A similar IgG deposition at the sarcolemma of skeletal muscles was also observed in 2-week-old *mdx* mice (**Figure 1A and fig. S1, A to D**), a widely-used DMD model which contained a point mutation in the exon 23 (*25*). Wild type littermates did not exhibit any immunoglobin deposition in their skeletal muscles (**Figure 1A and fig. S1, A to D**). Notably, at 2 weeks of age, neither *Dmd*^E4*^ nor *mdx* mice revealed signs of muscular damage and/or notable inflammation, consistent with previously report (*26*). Serum creatine kinase, a clinical biomarker of possible muscle damage, was comparable to the wild type littermates in 2-week-old *Dmd*^E4*^ or *mdx* mice (**Figure 1B**); infiltrating inflammatory cells and myofibers with centralized nuclei, a phenomenon associated with muscular regeneration, were not observed in skeletal muscles (**fig. S1, E and F**). Together, these observations suggest that IgG was ectopically deposited in the plasma membrane of dystrophin-deficient myofibers preceding most pathological abnormalities.

**Figure 1.**
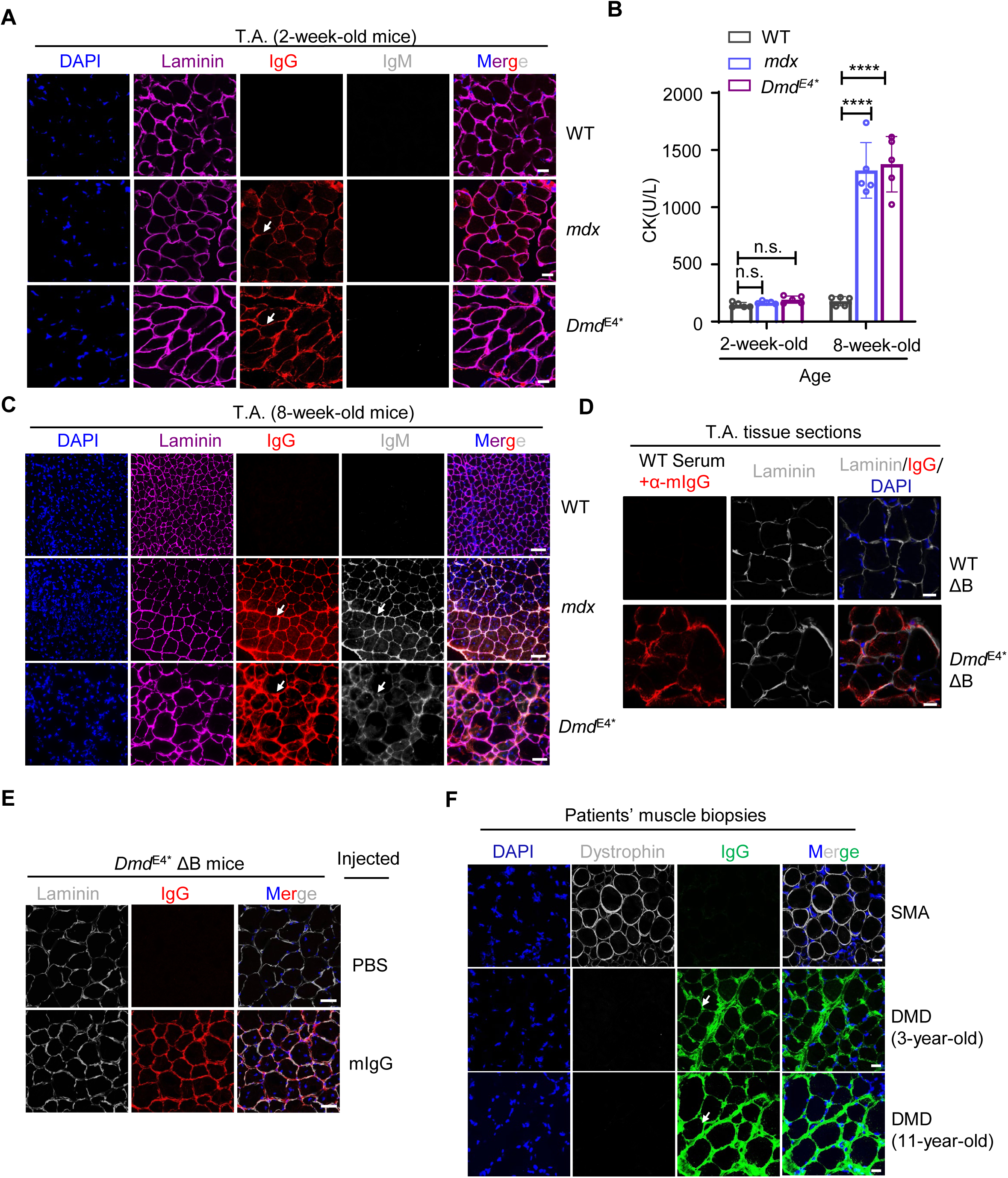
**Accumulation of non-autoreactive immunoglobins at the sarcolemma of dystrophin-deficient skeletal muscles** (**A**). Accumulation of IgG on the sarcolemma of tibialis anterior (T.A.) muscles in 2-week-old *Dmd*^E4*^ and *mdx* mice. Tissue sections (10 μm thick) were stained with Alexa Fluor 555-labeled anti-mouse IgG, Alexa Fluor 488-labeled anti-mouse IgM, Alexa Fluor 647-labeled anti-Laminin, and DAPI. Data are representative of five mice in each group. Scale bars represent 20 μm. Immunofluorescence staining of the heart, the rectus femoris (R.F.), and the diaphragm was included in **Fig. S1, B to D**. Arrows highlight the sarcolemma IgG staining. (**B**). Serum creatine kinase (CK) was determined in 2-week-old or 8-week-old *Dmd^E4*^* mice, *mdx* mice and littermate controls. Each dot represents one individual mouse (n=5). Error bars denote the standard deviation of the mean. ****, p<0.0001, n.s., not statistically significant based on Student’s *t* test. (**C**). Accumulation of IgG and IgM to sarcolemma of skeletal muscles in 8-week-old *Dmd^E4*^* and *mdx* mice. As in (**A**), T.A. muscles were stained with anti-mouse IgG, anti- mouse IgM, anti-Laminin, and DAPI. Scale bars represent 100 μm. Data are representative of six mice in each group. Arrows highlight the sarcolemma IgG and IgM staining. (**D**). Serum IgG of WT mice binds to DMD muscles *in vitro*. T.A. muscles from either uMT mice or uMT *Dmd*^E4*^ mice were stained with serum from WT mice, followed by Alexa Fluor 555 labeled anti-mouse IgG. Scale bars represent 20 μm. Data are representative of six mice in each group. (**E**). Purified IgG of WT mice bind to DMD muscles *in vivo*. IgGs (15 mg/kg) purified from WT mice were injected intraperitoneally into 1-week-old *Dmd*^E4*^ ΔB mice. T.A. muscles of the recipients were stained with Alexa Fluor 555 labeled anti-mouse IgG. Scale bars represent 50 μm. Data are representative of five mice in each group. (**F**). Accumulation of IgG at the sarcolemma in patients with DMD. Muscle biopsies from a 3-year-old DMD patient and a 11-year-old DMD patient were analyzed for dystrophin expression and IgG accumulation using immunofluorescence staining. Muscle biopsies from a patient with spinal muscular atrophy (SMA) were included as a control. Scale bars represent 20 μm. Arrows highlight the sarcolemma IgG staining.

This accumulation of antibodies was intensified with the progression of muscular dystrophy and/or the maturation of the immune system. In 8-week-old *Dmd*^E4*^ and *mdx* mice, both IgG and IgM were detected at the plasma membrane of cardiac and various skeletal muscle cells (**Figure 1C, Fig. S2A and Fig. S2, C to E**). IgG1, IgG2b, and IgG2c were the major isotypes, consistent with their dominance in the serum (**Fig. S2B**). Super resolution microscopic analysis further indicated that IgG mainly localized to the sarcolemma based on its proximity to laminin (**Fig. S2F**). Immunofluorescence analysis of the whole cross-sectional muscle biopsy revealed that majority of myofibers in the T.A. (**Fig. S3A and Movie S1**) and R.F (**Fig. S3B**) of *Dmd*^E4*^ mice exhibited IgG deposition. Importantly, the myofibers with IgG deposition were not associated with other serum proteins (*e.g.,* Albumin, **Fig. S2G**). No IgG and IgM accumulation was observed in the kidney or liver of the *Dmd*^E4*^ and *mdx* mice (**Fig. S3, C and D**).

IgG accumulation on the sarcolemma of dystrophin-deficient myofibers could be resulted from either increased autoreactive antibodies or ectopic expression of antibody-binding molecules (*e.g.,* Fc receptors). To test these two possibilities, we first utilized serum from WT, *Dmd*^E4*^ or *FasL* ^gld/gld^ mice to stain muscle tissues of uMT mice, which lack endogenous B cells and antibodies (*27*). As a positive control, serum from *FasL*^gld/gld^ mice (*28*), containing autoreactive antibodies similar to those in lupus, produced strong IgG and IgM staining signals. In contrast, serum from 2-week-old *Dmd*^E4*^ mice presented negative staining of either IgG or IgM in the muscle tissues of uMT mice (**Fig. S3E**), suggesting an absence of autoantibodies in the serum of *Dmd*^E4*^ mice at the early stage of the disease. On the other hand, IgG from WT mouse serum was able to stain sarcolemma of skeletal muscles from B cell-deficient *Dmd*^E4*^ mice (*Dmd*^E4*^ uMT mice, named as *Dmd*^E4*^ΔB mice thereafter), but not those from B cell-deficient WT mice (WT ΔB mice) (**Figure 1D**), suggesting dystrophin-deficiency is required for sarcolemma IgG deposition. Furthermore, when purified IgG from WT mice was transferred to *Dmd*^E4*^ΔB mice, the IgG accumulation was observed at the plasma membrane of skeletal muscles (**Figure 1E**) in a similar fashion as observed in B cell replete *Dmd*^E4*^ mice. Collectively, these *in vitro* and *in vivo* results demonstrate that non-autoreactive immunoglobulins are able to bind to dystrophin-deficient myofibers in the early stage of DMD disease.

In contrast to their sarcolemma deposition in young DMD mice, in 1-year-old *Dmd*^E4*^ mice, IgG and IgM were primarily observed in the cytoplasm of necrotic myofibers (**fig. S2H**), which was identified by RIPK3 aggregation as reported previously (*29*). While the intracytoplasmic accumulation of IgG at a later stage is likely a result of a passive influx of serum proteins into muscle fibers due to the leakage of the dystrophic myofibers (*30*), the sarcolemma IgG deposition prior to pathohistological changes in DMD has not been reported previously.

### Intramuscular IgG accumulation in patients with DMD

Consistent with observations in murine models, IgG accumulation was found in muscle biopsies from patients with DMD as young as three years old (**Figure 1F**), a stage when syndromes are often subtle(*31*). This IgG accumulation was consistently observed in patients with DMD up to 11 years old (**Figure 1F, Fig. S4A**), the oldest age we were able to obtain sample from. In addition to DMD, low levels of IgG accumulation in muscle biopsies were observed in patients with Becker muscular dystrophy (BMD), a genetic disorder caused by in-frame mutations in the DMD gene with milder symptoms(*32, 33*). The IgG accumulation in BMD patients appeared to be inversely correlated with dystrophin levels (**Fig. S4B**), though sample number was limited. Conversely, no or little IgG deposition was observed in muscle biopsies from patients with other forms of neuromuscular diseases, including SMA (**Figure 1F)**, Ataxia, Limb-girdle muscular dystrophy (type 4), and myotonic dystrophy (**Fig. S4B**). Taken together, these findings, along with the results from murine models, indicate that the loss of dystrophin leads to a characteristic IgG accumulation in myofibers of patients with DMD and BMD.

### IgG exacerbates DMD pathogenesis

We next investigated the roles of immunoglobins in the pathogenesis of DMD by comparing *Dmd*^E4*^ mice with *Dmd*^E4*^ ΔB mice. As expected, neither B cells nor antibodies were detectable in *Dmd*^E4*^ ΔB mice (**fig. S5, A and B**), thus IgM and IgG were absent in the muscle tissues (**fig. S5C**). Strikingly, the 8-week-old *Dmd^E4*^* ΔB mice exhibited substantially milder muscular dystrophy compared to their B cell-containing counterparts (**fig. S5, D to I**). Such improvement was sustained up to 12-month-old *Dmd^E4*^* ΔB mice, and the life span of *Dmd*^E4*^ mice was significantly extended when compared to *Dmd*^E4*^ mice (**Figure 2A**). Serum creatine kinase levels in 12-month-old *Dmd^E4*^* ΔB mice were reduced (from 1281.8±181.2U/L to 355.1±72.9U/L) to levels similar as in age-matched WT mice (**Figure 2B**). Accelerated muscle fatigue was largely rescued following repetitive measurement of muscle strength (**Figure 2C and fig. S5D**). Fibrosis in the various muscles (Masson’s staining) was greatly diminished in *Dmd*^E4*^ ΔB mice (**Figure 2, E and F**). Moreover, *Dmd*^E4*^ ΔB mice were protected from kyphosis (*34*), indicating improved strength in spine-supporting muscles (**Figure 2D**).

**Figure 2.**
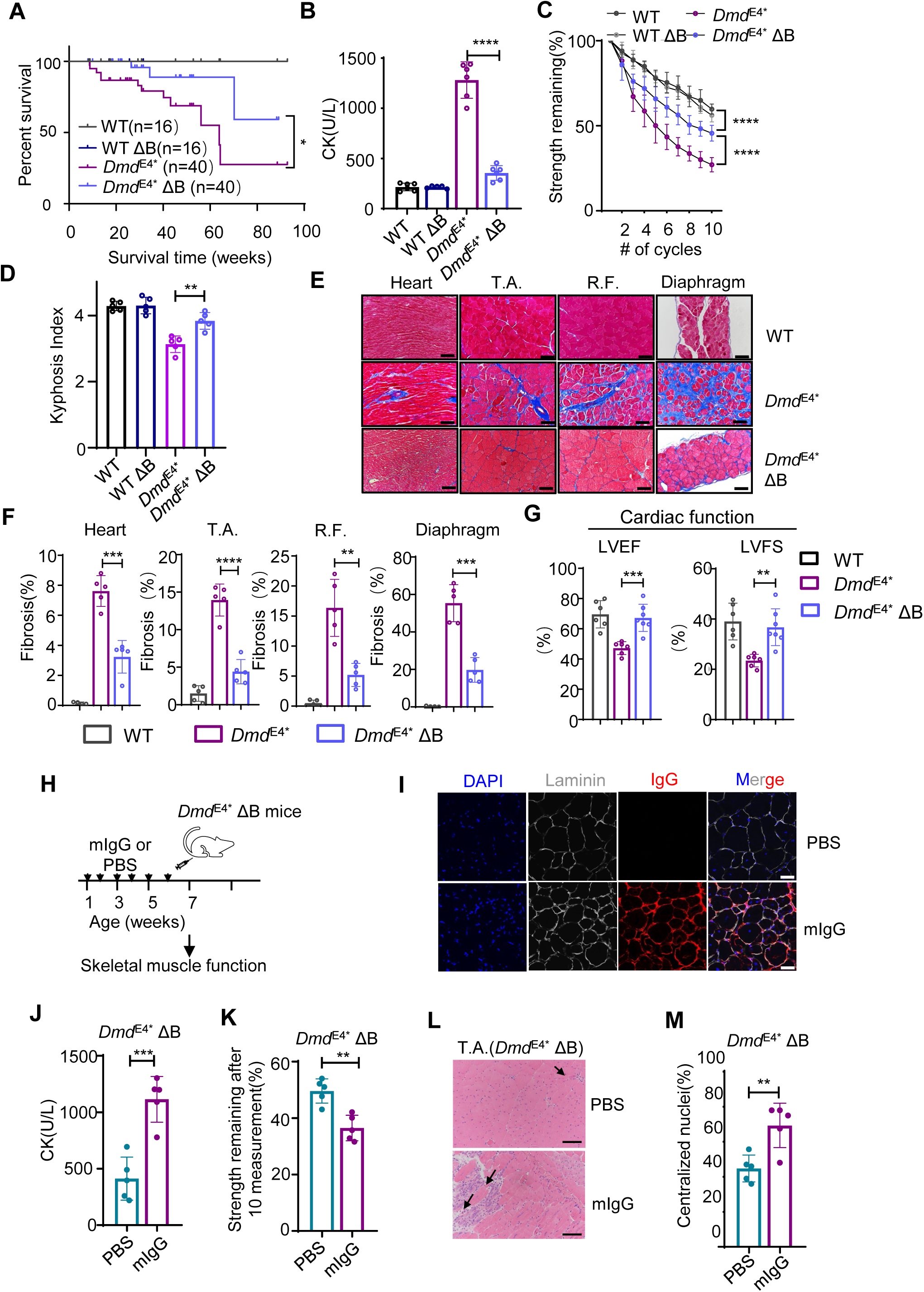
**Elimination of B cells improves muscle dystrophy in DMD mice** (**A**). Lack of B cells prolonged the life span of *Dmd^E4*^* mice. Kaplan-Meier survival curve analysis of *Dmd^E4*^* mice and *Dmd*^E4*^ ΔB mice. *, p=0.0278 based on the Mantel-Cox test. (**B**). Serum creatine kinase (CK) was determined in 12-month-old *Dmd*^E4*^ ΔB, *Dmd^E4*^* mice, and WT littermates. Data are a summary from at least 5 mice mice in each group. Each dot represents one individual mouse. (**C**). 12-month-old *Dmd*^E4*^ ΔB mice were protected against the decline of grip strength during a fatigue-inducing condition. 10 whole-body grip strength tests were conducted with a short interval (10 seconds) between each test, and the reduction in strength was calculated by normalized to the maximal grip strength. Data are summarized of 6 mice in each group. (**D**). 12-month-old *Dmd*^E4*^ ΔB mice were protected from kyphosis. Spinal curvature of 12-month-old WT, WTΔB, *Dmd^E4*^* and *Dmd^E4*^*ΔB mice was determined via µ-computed tomography (micro-CT) images as shown in **Fig.S5F**. Kyphosis indexes (KI) were calculated as reported (*34*). Briefly, A1 is determined as the distance between the posterior edge of the last cervical vertebra (C7) to the posterior edge of the sixth lumbar vertebra (L6). A2 is the longest distance from the dorsal border of the vertebral body to line A1. KI was defined as A1/A2. Data are a summary of five mice in each group. (**E**,**F**). Mason staining of various muscle tissues from *Dmd*^E4*^ ΔB, *Dmd^E4*^*, and WT littermate control mice at 12 months of age. T.A.: tibialis anterior, R.F.: rectus femoris. The data shown are either representative (**E**) or summary (**F**) of 5 mice in each group. Error bars stand for the standard deviation of the mean. Each dot represents one individual mouse. Scale bars represent 100 μm. (**G**). Lack of B cells alleviated cardiomyopathies in 12-month-old *Dmd*^E4*^ mice. Echocardiography analysis was performed in WT, *Dmd* ^E4*^ and *Dmd* ^E4*^ ΔB mice. LVEF (left ventricle ejection faction) and LVFS (left ventricle fractional shortening) were calculated based on the measurements collected from five consecutive cardiac cycles in the M-mode. Data are a summary of at least 6 mice in each group (Each dot represents one individual mouse). (**H**). Schematic showing transferring purified IgG into *Dmd*^E4*^ ΔB mice. Purified IgG (15mg/kg) from WT mice was injected intraperitoneally into 1-week-old *Dmd*^E4*^ ΔB mice twice a week for six weeks. Muscle functions were analyzed on the mice at the age of 7 weeks, and the results were compared to those of *Dmd*^E4*^ ΔB mice that received PBS injection (mock). (**I**). Purified IgGs bind to DMD muscles *in vivo*. Six weeks after the injection, T.A. muscles were stained with Alexa Fluor 555 labeled anti-mouse IgG, Alexa Fluor 647-labelled anti-mouse Laminin, and DAPI. Scale bars represent 50 μm. Data are representative of five mice in each group. (**J**). Transferring IgG from WT mice exaggerated muscular damages in *Dmd*^E4*^ ΔB mice. Six weeks after the first injection, levels of CK in the serum were determined. Data are a summary of five mice in each group. (**K**). Remaining grip strength after 10 consecutive measurement was compared between *Dmd*^E4*^ ΔB mice receiving IgG and their counterparts receiving PBS. Data are a summary of five mice in each group. (**L**). H&E staining of the T.A. muscle sections. Arrows indicate the area with inflammatory cell infiltration. Scale bars represent 100 μm. (**M**). Summary of myofibers with centralized nuclei in T.A. muscles. Data are a summary of five mice in each group. Each dot represents one individual mouse. Error bars stand for the standard deviation of the mean. ** p<0.01, *** p<0.001, **** p<0.0001 in two-tailed Student’s *t* test (**B**, **D, F, G, J, K, M**) or two way ANOVA test (**C**).

In addition to enhanced skeletal muscle function, dilated cardiomyopathy was largely attenuated in 12-month-old *Dmd*^E4*^ ΔB mice. These mice displayed improved left ventricular ejection fraction (LVEF%, increased from 47.2±4.3% to 67.2±9.0%) and left ventricular fractional shortening (LVFS%, increased from 23.5±2.5% to 36.7±7.3%) levels when compared to B cell replete *Dmd*^E4*^ mice (**Figure 2G**). These cardiac measurements were similar to those observed in age-matched wild-type mice (LVEF, 69.6±9.0%; LVFS, 39.0±7.3%) (**Figure 2G**). Similar milder phenotypes were observed in B cell-deficient *mdx* mice (*mdx* ΔB) (**fig. S6, A to F**).

In contrast, transferring purified IgG from wild-type mice to *Dmd*^E4*^ ΔB mice accelerated the progression of muscular dystrophy (**Figure 2H**). The purified IgG accumulated at the plasma membrane of skeletal muscles (**Figure 2I**)*. Dmd*^E4*^ ΔB mice receiving the IgG transfers presented elevated serum CK levels (from 411.7±191.2 U/L to 1114.4±202.7 U/L, **Figure 2J**), increased susceptibility to muscle fatigue upon repeated muscle strength measurements (**Figure 2K**), increased infiltrating inflammatory cells (**Figure 2L)**, and more myofibers with centralized nuclei (**Figure 2M**). Together, these findings suggest that B-cell produced IgGs accelerate DMD pathogenesis.

### Elevated FcγRI levels in *Dmd* myofibers promotes IgG accumulation

To understand how dystrophin-deficient myofibers recruit immunoglobins, we immunoprecipitated IgG and their associated proteins from muscle lysates of *Dmd*^E4*^ mice using Protein A/G beads. Muscle lysates from WT uMT mice and *Dmd*^E4*^ uMT mice, both lacking immunoglobins deposition in the muscles, served as negative controls. Immunoprecipitants were then analyzed with LC-MS/MS (**Figure 3A**). As expected, immunoglobulins and complement proteins were enriched in the immunoprecipitated fraction of *Dmd*^E4*^ mice compared to the controls. Notably, FcγRI (CD64), a high-affinity Fc receptor for IgGs(*35*), and Trim21, an intracellular Fc binding protein (*36*), were both highly enriched in the immunoprecipitated fraction from the *Dmd*^E4*^ mice compared to the controls (**Figure 3B and table S1**). The other Fc receptors (*e.g.* FcγRIIA, FcγRIIB (CD32)) were not identified (**table S1**). Indeed, immunofluorescence staining (**Figure 3C**) and RNA-scope analysis (**Figure 3, D and E and fig. S7C**) validated that FcγRI was markedly upregulated in skeletal muscles of the *Dmd*^E4*^ mice.

**Figure 3.**
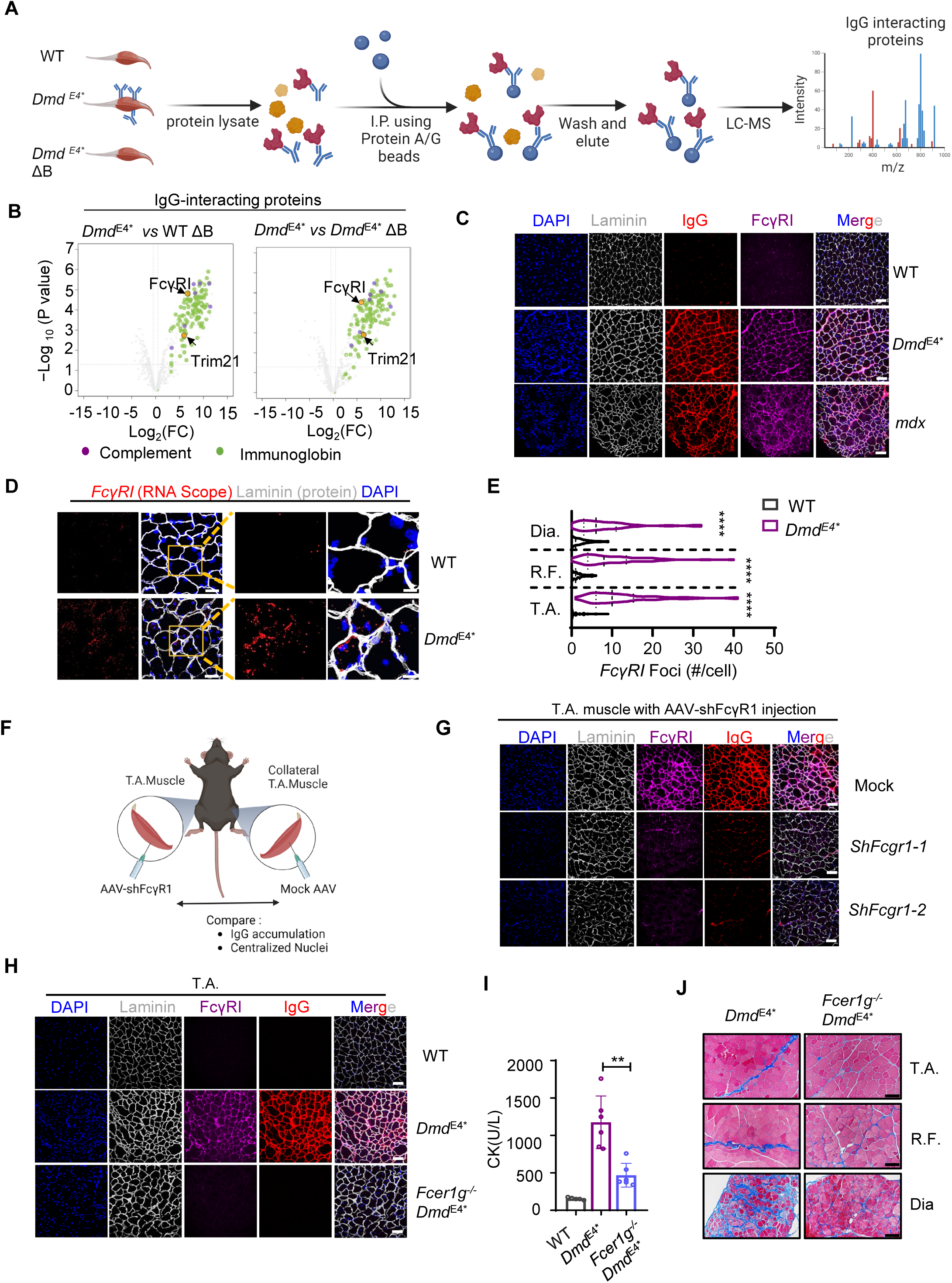
**FcγR1 recruits IgG to the dystrophin-deficient myofibers** (**A**). A workflow Schematic showing identification of IgG-Interacting Proteins via Immunoprecipitation and LC-MS/MS. Total tissue lysate of skeletal muscles from the *Dmd*^E4*^ mice was incubated with magnetic Protein A/G beads, and immunoprecipitates were analyzed using Mass spectrometry. Immunoprecipitates from WT ΔB mice and *Dmd*^E4*^ ΔB mice were included as negative controls. (**B**). Immunoglobin in the dystrophic muscles was associated with FcγR1.As in (**A**), enriched proteins were identified as p_adj_<0.001 and Log_2_ |FC|>=2. Immunoglobulins (green), complement proteins (purple), and known Fc receptors were indicated. Data are a summary of two independent experiments. (**C**). Ectopic presence of FcγR1 in DMD muscles. Tissue sections from 8-week-old *Dmd*^E4*^ and *mdx* mice were stained with Alexa Fluor 647-labelled anti-mouse FcγR1 and Alexa Fluor 555-labeled anti-mouse IgG. Laminin staining was included to define individual myofibers. Scale bars represent 100 μm. Data are representative of six mice in each group. **(D,E).** Elevated *FcγR1* transcript in DMD myofibers. As in (**D**), presence of FcγR1 in the T.A. was determined using RNAscope analysis. Data are representative of five mice in each group. Representative RNA-scope analysis of the rectus femoris (R.F.) and the diaphragm was included in **Fig. S7C.** Scale bars represent 50 μm (left panels) and 20 μm (right panels) respectively. (**E**). Summary of levels of *FcγR1* transcripts in T.A., R.F.,and diaphragm determined by RNA-scope. Each dot represents one individual myofiber. **(F,G).** Knockdown FcγR1 in myofibers diminished IgG accumulation in *Dmd*^E4*^ mice. (**F**). Schematic workflow: one side T.A. muscle of 4-week-old *Dmd*^E4*^ mice were injected with AAV9 virus particles containing shRNAs against Fcgr1 (sh-Fcgr1-1 or sh-Fcgr1-2). Four weeks after the treatment, levels of FcγR1 and accumulation of IgG were compared between the treated T.A. muscles and the counterpart control (mock AAV-treated T.A. muscle from the same animal) by immunofluorescence staining. (**G**). Representative immunofluorescence staining. Scale bars represent 100 μm. Summary of the result was included in the **Fig. S8A .** (**H**). Tissue sections from 6-week-old *Fcer1g^-/-^* (FcRγ-deficient) *Dmd*^E4*^ mice were stained with Alexa Fluor 647-labelled anti-mouse FcγR1 and Alexa Fluor 555-labelled anti-mouse IgG. Data are representative of five mice in each group. Scale bars represent 100 μm. (**I**). Genetic ablation of FcγR1 alleviates muscular dystrophy. 6-week-old *Fcer1g^-/-^ Dmd*^E4*^ mice and *Dmd*^E4*^ mice were analyzed for serum CK levels. Data are a summary of five or six mice in each group, and each dot represents one individual mouse. (**J**). Seven-month-old *Fcer1g^-/-^ Dmd*^E4*^ mice and *Dmd*^E4*^ mice were analyzed for fibrosis of various muscle tissues with Masson’s staining. Scale bars represent 100 μm. Data are reprenstative of five or six mice in each group, and summarized data are included in **Fig. S9D**.

FcγR1 is typically associated with myeloid cells such as macrophages, which are also the most abundant immune cells in DMD muscles. However, in *Dmd^E4*^TdTomato ^lox-STOP-lox^* Lyz-Cre mice where all the myeloid lineages and their derivatives were labelled with TdTomato. Immune fluorescence staining showed that majority (∼78%) of FcγR1 staining co-localized with Laminin rather than with TdTomato-positive cells in skeletal muscles (**fig. S7, A and B**). Moreover, elevated levels of FcγR1 were observed on myofibers in *Dmd*^E4*^ΔB mice, where B cells and antibodies are absent, although the levels were slightly reduced (**fig. S7D**). Similar to the observation in murine models, FcγRI was also detected in the muscle biopsy samples from patients with DMD, but not with Ataxia or myotonic dystrophy (**fig. S7E**). These findings indicate a non-canonical presence of FcγR1 in DMD muscle tissues, similar to the recent observation in joint sensory neurons in which FcγR1 contributes to arthritis pain (*37, 38*).

We then examined whether the elevated FcγRI was critical for intramuscular IgG accumulation and/or DMD progression. We first utilized adeno-associated virus vectors (AAV) 9 to deliver two independent shRNAs targeting FcγR1 (shFcgr1-1 or shFcgr1-2) to the T.A. via intramuscular injection (**Figure 3F**). AAV treatment led to over 80% reduction of FcγR1 on *Dmd*^E4*^ myofibers determined by immunofluorescence staining (**Figure 3G and fig. S8A**). Concomitant with the attenuated levels of FcγR1, both IgG deposition (**fig. S8A**) and muscular dystrophy as measured by cells with centralized nuclei (**fig. S8B**) were reduced in shRNA-treated myofibers compared to contralateral mock-treated muscles. Moreover, within the same T.A. muscle receiving the AAV injection, myofibers with higher AAV transduction (as indicated by GFP expression) exhibited lower levels of FcγR1 and less IgG accumulation than their un-transduced or less transduced counterparts (**fig. S8, C and D**). To further validate the relationship between ectopic FcγRI expression and intramuscular IgG accumulation, we crossed *Dmd*^E4*^ mice with Fc common γ chain (encoded by *Fcer1g*) KO mice (*39*) lacking surface expression of FcγRI, FcγRIII, and FcεRI receptors. In *ex vivo* primary myofiber culture, myofiber isolated from *Dmd*^E4*^ mice demonstrated elevated levels of FcγR1 and was able to bind to supplemented purified mouse IgG. In contrast, myofibers isolated for the Fcer1g^-/-^ *Dmd^E4*^* mice were unable to bind to supplemented IgG (**Fig. S.8, E-H**), indicating ectopic FcγRI in DMD deficient myofiber is accountable for IgG deposition. Furthermore, in the muscle tissue of *Fcer1g*^-/-^ *Dmd*^E4*^ mice, IgG deposition was not observed (**Figure 3H**). Importantly, *Fcer1g*^-/-^ *Dmd*^E4*^ mice exhibited milder muscular dystrophy phenotypes comparing to the *Dmd*^E4*^ mice, including reduced serum CK levels (from 1175.6±352.6U/L to 468.0±159.0U/L **Figure 3I**) and improved skeletal muscle functions (**fig. S9A**). Histological abnormalities were also alleviated, including inflammatory cell infiltration (**fig. S9B**), regenerating myofibers (**fig. S9C**), and fibrosis (**Figure 3J and fig. S9D**). These results together indicate a critical role of FcγR1 in recruiting IgG and promoting DMD pathogenesis.

### Loss of dystrophin elevates FcγR1 on myofibers

Since ectopic FcγR1 expression is observed in dystrophin deficient muscle tissue from mouse models and patients’ biopsies, we asked whether loss of dystrophin was causative to the ectopic expression of FcγR1. We restored a truncated but functional dystrophin in the myofibers of *Dmd*^E4*^ mice using AAV9 carrying a cytidine base editor and an sgRNA targeting 5’SS of the mutation-containing exon (exon 4). The intramuscular AAV treatment in 4-week-old *Dmd*^E4*^ mice induced ∼22% of exon skipping (**Figure 4, A and B**) and partially restored dystrophin (**Figure 4C**) as being reported previously (*12*). In response to the partial restoration of dystrophin 4 weeks after AAV treatment, a reduced level of FcγRI and IgG deposition was observed *via* immunofluorescence staining (**Figure 4, C and D),** compared to the mock-treated muscles on the opposite side of the same mice, suggesting loss of dystrophin can upregulate FcγRI in myofibers.

**Figure 4.**
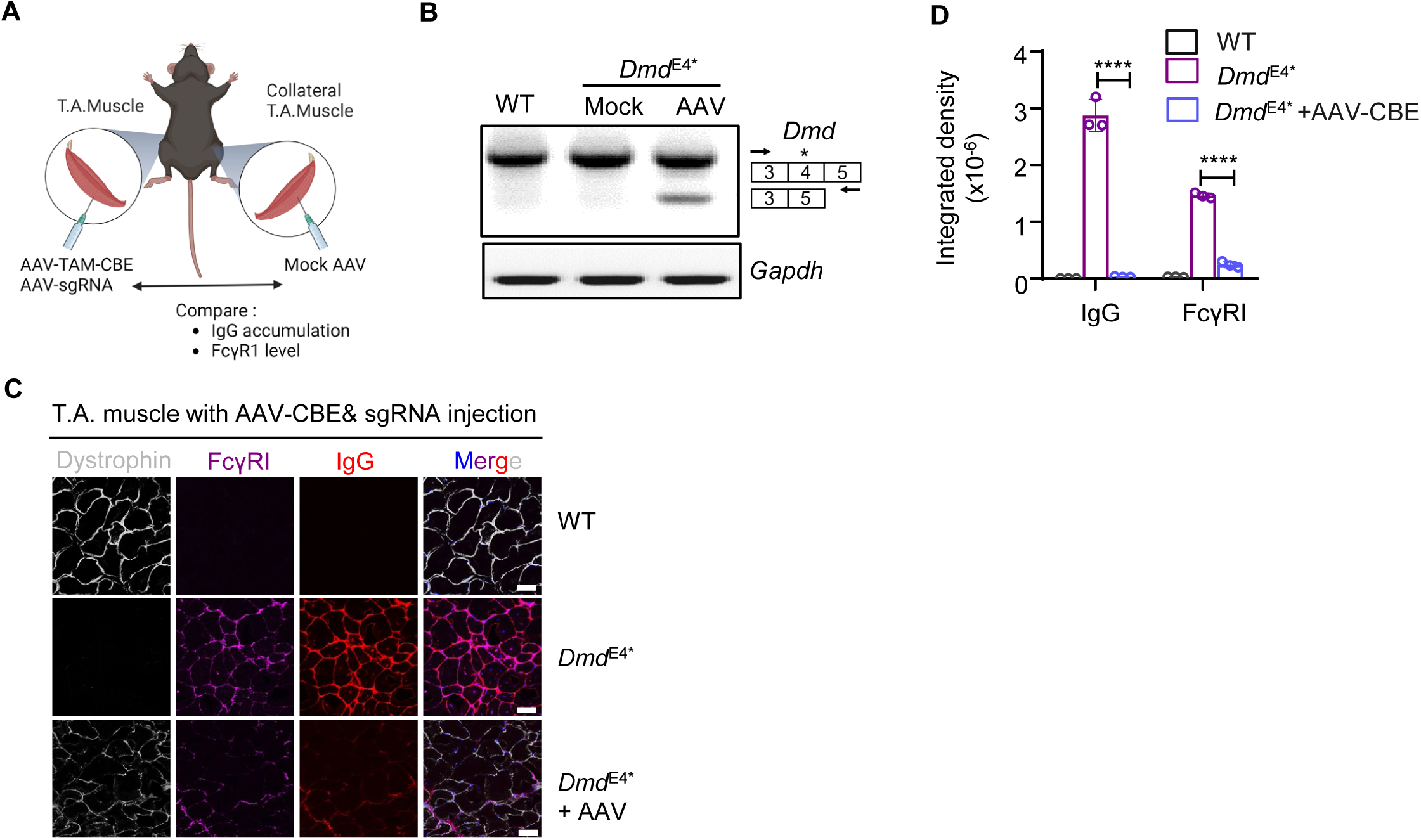
**Loss of dystrophin upregulates FcγR1** (**A-D**). Restoration of dystrophin reduced levels of FcγR1 and IgG accumulation. (**A**). A schematic overview. Right T.A. of 4-week-old D*md^E4*^* mice were injected with AAV9 containing the TAM-CBE base editor and sgRNA to induce exon skipping and restore dystrophin expression (2*10^12^ vg/T.A.). Three weeks after the injection, levels of FcγR1 and Dystrophin, and IgG accumulation were compared between the treated and the contralateral untreated T.A. of the same mouse. (**B**). The skipping of the mutation-containing exon (exon 4) was determined using RT-PCR. (**C,D**). Representative (C) or summary (D) of the immunofluorescence staining results. Integrated density of the staining was determined using Image J. Data are representative (**B,C**) or summary (**D**) of three mice in each group, and each dot represents one individual mouse. Scale bars represent 100 μm. Error bars stand for the standard deviation of the mean. * p<0.05, ** p<0.01, *** p<0.001 in two-tailed Student’s *t* test (**D**).

### Temporary depletion of B cells eliminates IgG accumulation and alleviates muscular dystrophy in two murine models of DMD

Having established that FcγRI and IgG accumulation in muscular tissues were critical early events of DMD pathogenesis, we investigated whether manipulation of B cells and antibody production could be exploited as a therapeutic strategy for DMD. Monoclonal antibodies against CD20 (*e.g.,* rituximab, ocrelizumab) have been successfully used in clinic to treat B cell malignancies and certain autoimmune diseases (e.g. multiple sclerosis) with limited side effects (*43*). We intraperitoneally injected 1-week-old *Dmd*^E4*^ mice with a B cell-depleting antibody specific to mouse CD20 (clone, MB20-11)(*44*) once a week for eight weeks (**Figure 5A**). The control mice were treated with a mouse IgG control antibody. Three weeks after the first anti-CD20 treatment, B cells were depleted by >95% in peripheral blood (**Figure 5B**). Correspondingly, systemic IgG and IgM levels in the serum were reduced by over 90% and 50%, respectively (**fig. S10, A and B**). As a result, IgG deposition in the muscular tissues of the anti-CD20 treated *Dmd*^E4*^ mice was greatly reduced as determined by immunoblotting (**fig. S10D**).

**Figure 5.**
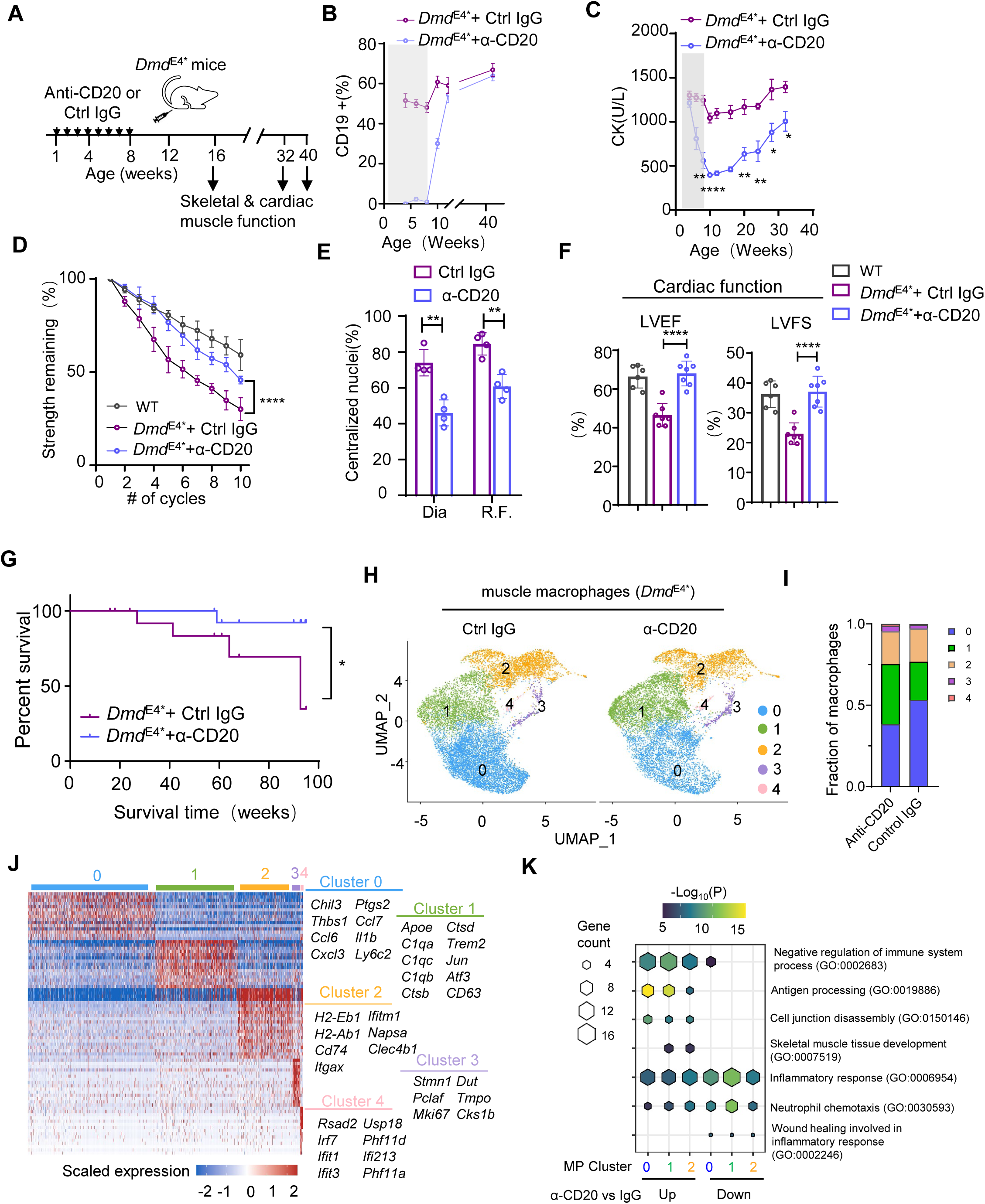
**Elimination of immunoglobins via B cell depletion alleviates muscular dystrophy in DMD mice** (**A**). Schematic of using anti-CD20 antibody to deplete B cells and eliminate antibody accumulation in DMD mice. Seven-day-old *Dmd* ^E4*^ mice were injected with anti-mouse CD20 (2 mg/kg) or control mouse IgG once a week for eight weeks. (**B**). As in (**A**), percentages of B cells (CD19^+^CD45^+^) in PBMC were determined using flow cytometry every week after the antibody treatment. Shaded area denotes the time periods during which mice received the antibody treatment. Data are a summary of six mice, and error bars stand for standard error of the mean. Note, the B cells quickly bounced back after the termination of the treatment. (**C**). As in (**A**), levels of serum CK were determined in the 4-week-old to 32-week-old *Dmd*^E4*^ mice, which received either control IgG or anti-CD20 treatment. The shaded area denotes the time periods during which mice received the antibody treatment. Data are a summary of three to six mice in each group, and error bars stand for standard error of the mean. (**D**). Anti-CD20 treatment prevented contraction-induced fatigue in *Dmd^E4*^* mice. Eight weeks after the last anti-CD20 treatment, the mice were tested for combined forelimb and hindlimb strength with a short interval (10 seconds) between each test. The reduction in strength was calculated by normalizing to the maximal grip strength after 10 grip tests. Data are summarized from 5 mice in each group. Error bars denote the standard deviation. **** p<0.0001 in two-way ANOVA test. (**E**). Summary of myofibers with centralized nuclei in R.F. and diaphragm from *Dmd*^E4*^ mice treated with anti-CD20 or control mouse IgG. Data are summarized from four mice in each group. (**F**). As in (**A**), echocardiograph analysis was performed in WT, *Dmd*^E4*^+control IgG and *Dmd*^E4*^ + anti-CD20 at 40 weeks of age. LVEF (left ventricle ejection faction) and LVFS (left ventricle fractional shortening) were calculated based on the measurements collected from three consecutive cardiac cycles in the M-mode. Data are a summary of six to seven mice in each group, and each dot represents one individual mouse. (**G**). Depleting B cells prolonged life span of the *Dmd^E4*^* mice. Kaplan-Meier survival curve analysis of *Dmd^E4*^* mice treated with either anti-CD20 (n=22) or control mouse IgG (n=18). p=0.0436 based on Mantel-Cox test. (**H-K**). Single cell RNA seq (scRNA-seq) analysis of skeletal muscle macrophages from anti-CD20 or control-IgG-treated *Dmd*^E4*^ mice. (**H**). Neonatal *Dmd*^E4*^ mice were treated with anti-CD20 or a control IgG as illustrated in Figure 5A. 12 weeks after the last injection, total CD45+ cells in skeletal muscles were sorted by flow cytometry and were analyzed by scRNA seq. Macrophages (Mono/Mac) cells were first identified as in **Fig.S12D** and subjected to UMAP analysis. 5 distinct clusters were identified and color coded. (**I**). Proportions of the five identified macrophage clusters, as shown in (**H**), were compared between control IgG-treated and anti-CD20-treated *Dmd*^E4*^ mice. (**J**). Heatmap of top 5-10 marker genes for each cluster is presented as normalized log2(counts) in all macrophages (in columns). Color-coded bars on the top indicate the clusters identified in (**H**). (**K**). Gene ontology analysis of significantly upregulated or downregulated genes between skeletal muscle macrophages from control IgG or anti-CD20-treated mice. Size of the hexagons represents the number of genes per GO term and the density represents the p value of enrichment after -log10 transformation. Color-coded numbers at the bottom indicate clusters identified in (**H**). Error bars stand for the standard deviation of the mean. * p<0.05, ** p<0.01, *** p<0.001, **** p<0.0001 in two-tailed Student’s *t* test (**C**, **E, F**) or two way ANOVA test (**D**).

Importantly, the anti-CD20-treated *Dmd*^E4*^ mice displayed significant phenotypic improvement compared to the control antibody-treated mice. Levels of CK in the serum were reduced by more than 50% eight weeks after the anti-CD20 treatment compared to control IgG-treated mice (from 1242.7±137.2U/L to 557.9±223.9U/L, **Figure 5C**). Functionally, the accelerated muscle fatigue following repetitive measurement of muscle strength (**Figure 5D**) and kyphosis (**fig. S10E)** were largely rescued. Histological analysis revealed decreased inflammatory cell infiltration (**fig. S10F**), reduced fibrosis (**fig. S10, G and H**), and fewer myofibers with central nuclei (**Figure 5E**) in the anti-CD20 treated mice compared to the control IgG-treated mice.

Notably, a transient depletion of B cells for eight weeks led to a prolonged benefit in the pathophysiological progression of DMD. Percentages of B cells and levels of IgM (**Figure 5B, fig. S10A)** in the blood were fully recovered four weeks after the last anti-CD20 treatment, yet levels of IgG remained at low levels for at least 16 weeks after the last treatment (135.5±40.8 ug/ml vs 548.1±183.2 ug/ml, **fig. S10B**). Six months after the anti-CD20 first treatment, *Dmd*^E4*^ mice still maintained low levels of serum CK (from 1176.6±73.6U/L prior to the treatment to 665.2±263.5U/L six months post treatment, **Figure 5C**) and is resistant to muscle fatigue (**fig. S10C**). Eight months after the treatment, cardiac function in the anti-CD20 treated group (LVEF, 68±6.5%;LVFS, 37±5.2%) remained comparable to the WT group (LVEF, 66.4±5.9%;LVFS, 36.2±4.5%), in contrast to the *Dmd*^E4*^ mice received control IgG treatment (LVEF%, 46.5±6% ; LVFS%, 23±3.6%), **Figure 5F)**. Ultimately, the lifespan of the *Dmd*^E4*^ mice was significantly extended in the anti-CD20 treated group compared to the IgG control group (**Figure 5G**).

Transient depletion of B cells using the anti-CD20 antibody also ameliorated muscular dystrophy in adult mice with established DMD. 8-week-old *Dmd*^E4*^ mice were treated with anti-CD20 once a week for eight weeks (**fig.**

**S11A**), resulting in a loss of B cells (**fig. S11B**). Although B cells bounced back two weeks after the last treatment, the anti-CD20 treatment afforded a lasting benefit to curtail dystrophy progression. Serum CK levels remained low (**fig. S11C**) and resistance to muscle fatigue (**fig. S11D**) were improved for at least 12 weeks following the last antibody treatment, at which time B cells were completely recovered to the prior levels (**fig. S11B**). Similar to the *Dmd*^E4*^ mice, *mdx* mice receiving anti-CD20 antibody (**fig. S11E**) exhibited a sustained reduction of muscular dystrophy, including reduced serum CK levels (**fig. S11F**) and resistance to muscle fatigue (**fig. S11G**). Together, these data demonstrate that B cell depletion was effective to both prevent and mitigate muscular dystrophy in DMD murine models.

### Antibody elimination shifts the inflammatory milieu in DMD muscles towards a pro-healing state

Antibody and immune complex deposition in tissues are known to trigger a cascade of inflammatory events in conditions such as glomerulonephritis and autoimmune lung disease. These events include the release of pro-inflammatory factors, activation of complement pathways, recruitment of pro-inflammatory macrophages, and inhibition of anti-inflammatory macrophages and immune factors(*45–47*). To elucidate how elimination of antibody accumulation in muscle tissue led to sustained improvement in muscular dystrophy, we analyzed the downstream inflammatory cascades affected by B cell depletion.

We found B cell depletion using the anti-CD20 treatment modified cytokine milieu in dystrophic muscles. OLINK proteomic analysis, which employs proximity extension assay (PEA), of interstitial fluid from *Dmd*^E4*^ muscles revealed elevated levels of IL1β, Ccl2, Ccl3, and CXCl1 when compared to wild-type controls (**fig. S12H**). Conversely, IL-10, an anti-inflammatory cytokine known to alleviate muscular dystrophy was reduced in DMD muscles (**fig. S12H**). Importantly, anti-CD20 treatment led to decreased Ccl2 levels and a concomitant increase in IL-10 (**fig. S12, I and J**) in the muscle of *Dmd*^E4*^ mice compared to the control IgG-treated mice.

Furthermore, the anti-CD20 treatment reduced the number of infiltrating leukocytes in the skeletal muscles of *Dmd*^E4*^ mice compared to the control IgG-treated mice (from 8.3×10^4^/g to 3.4×10^4^/g, **fig. S12C**) even after peripheral B cells were fully recovered. Within all the infiltrating leukocytes, the numbers of macrophages were dramatically reduced following the anti-CD20 treatment (from ∼5×10^4^/g to ∼2×10^4^/g, **fig. S12, A and C**), while B cells, T cells, and Treg cells were not altered (**table S2**). Notably, the ratio of CD86-expressing proinflammatory macrophages decreased, while CD206-expressing anti-inflammatory macrophages increased in the anti-CD20-treated mice compared to control IgG-treated mice (**Fig. S12, B and C**).

We further isolated CD45^+^ leukocytes from the skeletal muscles and conducted single-cell RNA sequencing (scRNA seq) analysis 12 weeks after the last treatment. Unsupervised clustering and UMAP analysis revealed 11 distinct clusters, with the majority of infiltrating leukocytes being mono/macrophages (Cluster 0,1,2, and 3)(**fig. S12, D and E**). We re-clustered the mono/macrophages into five distinct clusters (MP cluster 0-4, **Figure 5H**). MP cluster 0, displayed high expression levels of proinflammatory macrophage-associated genes (Thbs1, Ccl6, IL1b, Ccl2) and monocyte markers (Ly6c2, Chil3), suggesting they are monocyte-derived proinflammatory macrophages (**Figure 5J and fig. S12F**). MP cluster 1 exhibited high expression of C1q genes (C1qa, C1qb, C1qc) (*48, 49*), cathepsins (Ctsb, Ctsd, Ctss) (*50*), TREM-2, and apolipoprotein E (Apoe), (**Figure 5J and fig. S12F**), resembling C1q macrophages, a subset of tolerogenic and immunosuppressive macrophages in tissue damage and tumors (*51–54*). MP cluster 2 and MP cluster 3 correspond to dendritic cells and proliferating macrophages respectively (**Figure 5J**). MP cluster 4 was highly enriched for IFN-inducible genes (Ifit2, Isg15, and Irf7), indicating they were IFN-induced macrophages.

Consistent with our flow cytometric analysis, ratios of proinflammatory macrophages (MP cluster 0) and anti-inflammatory C1q macrophages (MP cluster 1) inverted reciprocally after the anti-CD20 treatment, while other clusters remained largely unchanged. Proinflammatory macrophages were decreased in anti-CD20-treated mice (from 53.2% to 38.4%) (**Figure 5, H and I**), while C1q anti-inflammatory macrophages were expanded (from 23.5% to 37.0%). Further gene ontology analysis showed that genes involved in negative regulation of inflammatory responses (GO:0002683) and cell junction disassembly (GO:0150146) were upregulated in all 5 clusters of macrophages from the anti-CD20-treated mice (*e.g.*C1qa, C1qb, and C1qc) (**Figure 5K and fig. S12G**). Conversely, genes involved in proinflammatory responses (GO:0006954) and neutrophil chemotaxis (GO:0030593) were suppressed in macrophages following the anti-CD20 treatment (**Figure 5K and fig. S12G**). Together our findings that the temporal B cell depletion via anti-CD20 treatment was able to rebalance pro- and anti-inflammatory responses in the dystrophic muscle, leading to delayed and mitigated DMD progression.

## Discussion

Recent therapeutic strategies for Duchenne muscular dystrophy (DMD) have primarily focused on two approaches: restoring dystrophin expression and targeting downstream pathological changes (*16*). While significant progress has been made in dystrophin restoration, this approach faces many challenges, including delivery issues, incomplete understanding of toxicity(*60–62*), limited efficacy, small patient segments, and high costs(*63, 64*). Our research unveils a novel mechanism in DMD pathogenesis that offers a promising alternative therapeutic avenue.

Our results show that the aberrant localization of FcγRI in dystrophin-deficient myofibers leads to immunoglobulin and immune complex accumulation in the muscle. The increased levels of FcγRI in dystrophic myofibers was supported by our findings in proteomic analysis of IgG-interacting proteins, immunofluorescence staining, RNA scope analysis, lineage tracing, and single myofiber culture. The critical roles of FcγRI in IgG accumulation and DMD pathogenesis are further corroborated by both genetic knockout and local shRNA knockdown experiments. Importantly, the FcγRI-dependent IgG accumulation precedes most pathological changes and likely initiates chronic inflammation together with other downstream events of dystrophin deficiency. Circulating immune complexes, formed under physiological conditions by multivalent antigens, immunoglobulins, and complement proteins, likely play a crucial role in this process (*65*). Similar to antibody and immune complex-mediated lung or renal injuries, IgG and immune complex deposition in dystrophin-deficient myofibers likely creates binding sites for complement proteins and Fc receptors on other immune cells, culminating in extensive muscle damage and loss of function.

Our research primarily focused on the accumulation of IgG at the sarcolemma as an initial event in the pathogenesis of DMD. This is in contrast to previous studies that have reported instances of intracellular IgG accumulation (as well as IgM and other serum proteins) in dystrophic muscles(*66*), largely linking it with membrane leakage and necrotic muscle fibers during the late stages of DMD pathogenesis. However, the binding of IgG to the plasma membrane of myofibers could potentially trigger downstream signaling events of FcγR1, which could directly lead to necrosis and subsequent intracellular accumulation of IgG/IgM in the later stages.. Nonetheless, necrosis, as a late-stage event, is likely the cumulative result of various factors and offers limited opportunities for intervention.

Notably, temporary B cell depletion alleviated muscular dystrophy progression for an extended period in both murine models. This suggests that transient B cell depletion may “reset” the immune microenvironment within DMD muscles, including a rebalancing of macrophage subtypes from pro-inflammatory to anti-inflammatory and/or pro-healing. This repositioning may halt the continuous cycle of muscle damage and repair in dystrophin-deficient muscles, providing therapeutic benefits.

Following anti-CD20-mediated B cell depletion in DMD mice, we observed significant changes in the immune milieu. The most notable alterations were the expansion of C1q macrophages (*51, 67, 68*) and a shift in muscle cytokine/chemokine profiles. Specially, we observed reduction of Ccl2 and Ccl3, which strongly attract monocytes and macrophages, and CXCL1, a potent neutrophil attractant. Conversely, levels of IL-10 were elevated in dystrophic muscles following the B cell depletion. Previous research has demonstrated that various cytokines, chemokines, and growth factors in dystrophic muscle can either promote inflammation-induced injury (e.g., Ccl2(*69*)) or facilitate regeneration (*e.g.* IL-10(*70*)). Moreover, immune cells such as macrophages may exhibit dual roles in DMD pathogenesis via transitioning between pro-inflammatory and anti-inflammatory phenotypes (*41, 71, 72*). These findings suggest a complex inflammatory response in the muscle of DMD, with IgG deposition emerging as a key early event initiating this intricate network. The protective effects of B cell depletion on disease progression corroborate this notion, implying the convergence of multiple downstream effector mechanisms in muscle damage.

Interestingly, B cell depletion in younger animals demonstrated better outcomes than in older ones. This could be attributed to prolonged antibody reduction or less severe pre-treatment muscle damage in younger subjects. These findings underscore the importance of early intervention and highlight the need for further research to optimize treatment protocols.

While B cell depletion shows great potential as a treatment strategy for DMD, further investigation is crucial to fully assess its clinical potential. This includes evaluating the long-term effects and safety of B cell depletion in DMD. Additionally, optimizing treatment protocols, including timing, duration, and frequency of B cell depletion, will be essential for maximizing therapeutic efficacy. Exploring potential synergies with other therapeutic approaches such as gene therapy or exon skipping could lead to more comprehensive treatment strategies. Furthermore, investigating alternative B cell depletion agents, such as antibodies targeting CD19, CD22, or BAFF, and CAR-T cells targeting CD19, may provide additional options for tailoring treatments to individual patient needs.

## ACKNOWLEDGMENT

We thank Dr. Zuoxian Meng, Heping Xu, Yichuan Xiao, Zhizhang Wang, and Yuan Chang for sharing critical reagents and providing suggestions. We thank advanced biomedical technology core facility, supercomputing center, Instrumentation and Service Center for Molecular Sciences and laboratory animal resource center at Westlake University for the facility support and technical assistance.

## Author contributions

Conceptualization: X.C.; Y.C.; Investigation: L.Z., W.G., M.H.,Z.L., E.Z., C.C., W.S., J.L., J.L., H.Q., L.Z., S.R., R.B. and J.W. ; Software: G.L; Formal analysis: L.Z., G.L., Y.G., and X.C.; Methodology: L.Z.; Resources: X.G., Z.W., Y.D., J.H., F.L., and W.J.; Writing – original draft: L.Z., W.G., Y.S., Y.C., and X.C.; Writing – review &editing: Y.C and X.C.; Supervision: X.C.; Visualization: L.Z., Y.C, and X.C.; Funding acquisition: Y.C and X.C.

## Data and materials availability

ScRNA seq data have been deposited in SRA under accession number PRJNA1023707. The MS data have been deposited in the ProteomeXchange proteomics Consortium (http://proteomecentral.proteomexchange.org) via the PRIDE partner repository with the following accession number: PXD052136 (Reviewer Username, Password: YEvJngHY). All other data are available in the manuscript or the supplementary materials.

## Supplementary Figures

Fig. S1. Accumulation of IgG in various skeletal muscles of 2-week-old *Dmd*^E4*^ and *mdx* mice

(A). Tissue lysate from 2-week-old *mdx*, *Dmd*^E4*^, and WT mice was analyzed by immunoblotting for IgM and IgG. GAPDH was included as loading controls. Data are representative of six mice in each group.

(**B-D**). Tissue sections of rectus femoris (R.F.) (**B**), diaphragm (**C**), and heart (**D**) (10 μm thick) were stained with Alexa Fluor 555-labeled anti-mouse IgG, Alexa Fluor 647-labelled anti-Laminin, Alexa Fluor 488-labeled anti-mouse IgM, and DAPI. Data are representative of five mice in each group. Scale bars represent 50 μm.

(**E-F**). There were no detectable inflammatory cell infiltration and muscles with centralized nuclei in 2-week-old *mdx* and *Dmd*^E4*^ mice. (**E**). Tissue sections of T.A. and R.F. muscles were examined with H&E staining. Scale bars represent 100 μm. (**F**). Summary of Myofibers with centralized nuclei in T.A. and R.F. muscles. Data are summarized from five mice in each group.

**Fig. S2. Accumulation of immunoglobins in skeletal and cardiac muscles of 8-week-old DMD mice**

(A). Immunoblot analysis of IgG accumulation. Total tissue lysate from the heart (left panel) or T.A. (right panel) was analyzed for IgM, IgG heavy chain (IgG H), or IgG light chain (IgG L). Data are representative of six mice in each group.

(B). Levels of IgG1, IgG2b, IgG2c, and IgG3 were analyzed in tissue lysate of T.A. muscles as in (**A**). Data are representative of four to six mice in each group. GAPDH was included as a loading control, and a molecular weight standard was marked on the left.

(**C-E**). Tissue sections of the heart (**C**), diaphragm (**D**), and R.F. (**E**) were stained with Alexa Fluor 555-labeled anti-mouse IgG, Alexa Fluor 647-labeled anti-Laminin, Alexa Fluor 488-labeled anti-mouse IgM. Data are representative of five mice in each group. Scale bars represent 50 μm.

(**F**). Super-resolution microscopy imaging of skeletal muscle tissues. The sarcolemma was stained with Laminin (gray) and indicated by arrows. The endomysial space, which represents the interstitial space between adjacent sarcolemmas, is highlighted by asterisks. Data are representative of four mice.

(**G**) Lack of albumin accumulation in the 8-week-old DMD muscles. T.A. muscles from 8-week-old *Dmd*^E4*^ mice were analyzed for the presence of albumin and IgG by immunofluorescence staining. Data are representative of five mice in each group. Scale bars represent 100 μm.

(**H**). Most myofibers with IgG accumulation did not show RIPK3 aggregation in 8-week-old *Dmd^E4*^* mice. Immunofluorescence staining showed the subcellular localization of RIPK3 in the T.A. of WT mice, 8-week-old *Dmd*^E4*^ mice, and 1-year-old *Dmd*^E4*^ mice. Notably, only in the myofibers of 1-year-old *Dmd*^E4*^ mice did the distribution of RIPK3 change to a more punctate and aggregated distribution, indicating the formation of necrosomes. Note, myofibers with RIPK3 punctate also had cytoplasmic IgM and IgG accumulation. Data are representative of three mice in each group. Scale bars represent 50 μm.

**Fig. S3. Characterization of antibody accumulation in the 8-week-old *Dmd* mice**

(**A,B**). Mosaic micrographs of the entire cross section of T.A. (**A**) or R.F. (**B**) from 8-week-old *Dmd*^E4*^ mice and WT mice. Data are representative of five mice in each group.

(**C,D**). Lack of antibody accumulation in non-muscular tissues of *mdx* or *Dmd*^E4*^ mice. Liver (**C**) and kidney (**D**) sections from 8-week-old WT, *mdx*, and *Dmd*^E4*^ mice were analyzed for the presence of IgG and IgM by immunofluorescence staining. Data are representative of five mice in each group. Scale bars represent 100 μm.

(**E**). Serum from 2-week-old *Dmd*^E4*^ mice or WT littermates were used as the primary antibody to stain T.A. tissue sections (10 μm thick) from WT μMT mice, which was followed by staining with Alexa Fluor 555-labeled anti-mouse IgG and Alex Fluor 488-labeled anti-mouse IgM. Data are representative of five mice in each group. Serum from lupus mice (*FasL* ^gld/gld^) was included as positive controls. Scale bars represent 100 μm.

**Fig. S4. IgG accumulation at sarcolemma of patients with Duchenne and Becker Muscular Dystrophy**

(A). IgG accumulation in patients with DMD. Muscle biopsies from patients with DMD at various ages were examined for the expression of dystrophin and the accumulation of IgG and IgM. The ages and DMD mutations of the patients were indicated on the right and left respectively. Tissue sections (10 μm thick) were stained with Alexa Fluor 488-labeled anti-human IgG, Alexa Fluor 594-labelled anti-Dystrophin, Alexa Fluor 647-labeled anti-human IgM, and DAPI. Each slide represents one patient. Scale bars represent 20 μm. N.A., genetic information of the patients was not available due to privacy regulations or because the patients were diagnosed before the widespread adoption of genetic testing.

(B). IgG accumulation in patients with BMD. Muscle biopsies from patients with BMD were analyzed for the expression of Dystrophin and the accumulation of IgG and IgM. Note a significant accumulation of IgG was observed in a patient with severe BMD with greatly reduced Dystrophin. Muscle biopsies from patients with Ataxia, Limb-girdle muscular dystrophy (type 4), and myotonic dystrophy were also included in the analysis. Tissue sections (10 μm thick) were stained with Alexa Fluor 488-labeled anti-human IgG, Alexa Fluor 594-labelled anti-Dystrophin, Alexa Fluor 647-labeled anti-human IgM, and DAPI. Scale bars represent 20 μm.

**Fig. S5. Improvement of muscular dystrophy in *Dmd*^E4*^ ΔB mice**

(A). Loss of B cells in the *Dmd*^E4*^ ΔB mice. Percentages of B cells (CD19^+^CD45^+^) in PBMC were determined using flow cytometry. Data are either representative (left) or summary (right) of six mice in each group.

(B). Loss of B cells, IgG and IgM in the *Dmd*^E4*^ ΔB mice. Serum IgG and IgM levels were determined using enzyme-linked immunosorbent assay (ELISA). Data are summary of six mice in each group.

(C). Lack of antibody accumulation in the skeletal muscles of *Dmd*^E4*^ ΔB mice. Tissue lysate from T.A. muscles of WT, *Dmd*^E4*^ and *Dmd*^E4*^ ΔB mice were analyzed by immunoblotting for IgM and IgG. Data are representative of five mice in each group.

(D). Enhanced skeletal muscle functions in 2-month-old *Dmd*^E4*^ ΔB mice. 10 whole-body grip strength tests were conducted with a short interval (10 seconds) between each test, and the reduction in strength was calculated by normalized to the maximal grip strength. Data are summarized from five mice in each group.

(E). Serum CK levels in 2-month-old WT, WT ΔB, *Dmd*^E4*^, and *Dmd*^E4*^ ΔB mice were determined. Data are summary of at least seven mice in each group. Each dot represents one individual mouse.

(F). Representative micro-CT images showing spinal curvature of 8-week-old WT, WT ΔB, *Dmd*^E4*^, and *Dmd*^E4*^ ΔB mice. Data are representative of four mice in each group.

(G). As illustrated in (**F**), Kyphosis indexes (KI) were calculated as reported (*34*). Each dot represents one individual mouse (n=4 in each group).

(**H**) Hematoxylin and eosin (H&E) staining of various muscle tissues from *Dmd*^E4*^ ΔB and *Dmd*^E4*^ mice at 8 weeks of age. T.A.: tibialis anterior, R.F.: rectus femoris. Arrows indicate areas with intensive immune cell infiltration. Scale bars: 100 μm. Data are representative of five mice in each group.

(**I**). Percentages of myofibers with centralized nuclei were calculated from 8-week-old WT, *Dmd*^E4*^ and *Dmd*^E4*^ΔB mice. Data are a summary of five mice in each group.

Error bars stand for the standard deviation of the mean. *** p<0.001, **** p<0.0001 in two-tailed Student’s *t* test (**B, E**, **G, I**) or two-way ANOVA test (**D**).

**Fig. S6. Improvement of muscular dystrophy in *mdx* ΔB mice**

(A). Improved skeletal muscle functions in 2-month-old *mdx* ΔB mice. 10 whole-body grip strength tests were conducted with a short interval (10 seconds) between each test, and the reduction in strength was calculated by normalized to the maximal grip strength. Data are summarized from 5 mice in each group. **** p<0.0001 in two-way comparison ANOVA test.

(B). Hematoxylin and eosin (H&E) staining of various muscle tissues from *mdx* ΔB and *mdx* mice at 12 months of age. T.A.: tibialis anterior, R.F.: rectus femoris. Data are representative of four to five mice in each group. Scale bars: 100 μm.

(C). Serum CK levels in 12-month-old *mdx* ΔB mice were compared to those in age-matched WT and *mdx* mice. Data are summarized from 5 mice in each group.

(D). Percentages of myofibers with centralized nuclei were determined in different skeletal muscles of 12-month-old WT, untreated *mdx* and *mdx* ΔB mice. Data are a summary from four or five mice in each group. Each dot represents one individual mouse.

(**E,F**). Masson’s staining of various muscle tissues from *mdx* ΔB, *mdx* and littermate control mice (WT) at 12-month-old -weeks of age. Data are representatives (**E**) or summary (**F**) of four to five mice in each group. Each dot represents one individual mouse. Scale bars: 100 μm.

Error bars stand for the standard deviation of the mean. * p<0.05, ** p<0.01, *** p<0.001, **** p<0.0001 in two-tailed Student’s *t* test (**C**, **D, F**) or multiple comparison ANOVA test (**A**).

**Fig. S7. Ectopic presence of FcγR1 on myofibers recruits IgG to dystrophic myofibers.**

(**A,B**). T.A. muscle from Lyz-Cre TdTomato ^LSL^ *Dmd*^E4*^ mice was analyzed for levels of FcγR1 and TdTommato by immunofluorescence staining. WT Lyz-Cre TdTomato ^LSL^ mice were included as controls. Arrows indicate Tdtommato^+^ cells, which also showed strong FcγR1 expression. Data are representative (**A**) or summary (**B**) of five mice in each group. Note, most FcγR1 signaling was derived from laminin-expressing cells, rather than TdTomato+ cells. Scale bars represent 50 μm.

(**C**). Representative RNA scope analysis of FcγR1 mRNA in R.F and Diaphram. Data are representative of five mice in each group. Scale bars represent 50 μm (left panels) and 20 μm (right panels) respectively.

(**D**). Expression of FcγR1 was sustained in the myofibers of *Dmd*^E4*^ ΔB mice, albeit with a minor reduction. T.A. muscle from eight-week-old *Dmd*^E4*^ΔB mice was analyzed for FcγR1 with immunofluorescence staining. Age matched WT mice and *Dmd*^E4*^ were included as controls. Data are representative or summary of four mice in each group.

(**E**). Elevated FcγR1 expression on myofibers from patients with DMD. Muscle biopsies from patients with DMD were examined for the expression of FcγR1 with immunofluorescence staining. Scale bars represent 20 μm.

**Fig. S8. Ectopic presence of FcγR1 on myofibers recruits IgG to dystrophic myofibers.**

(**A-B**). As in **Figure 3G**, levels of FcγR1 and IgG accumulation of myofibers were determined using Image J (**A**). Myofiber damage was determined by identifying myofibers with centralized nuclei (**B**). Scale bars represent 100 μm. Data are summary of four to six mice in each group, and each dot represents one individual mouse.

(**C,D**). As in **Figure 3F**, the T.A. muscle injected with AAV containing sh-Fcgr1 and GFP was analyzed by immunofluorescence staining. GFP ^hi^ and GFP ^low^ myofibers were identified, and levels of IgG accumulation and FcγR1 were analyzed. (**C**). High-magnification view of the circled GFP ^hi^ and GFP ^low^ myofibers.(**D**). As in (**A**), quantitative analysis of IgG and FcγRI staining intensities in the top 20% (GFP^hi^) and bottom 20% (GFP^low^) of myofibers.

(**E-H**). Myofibers were isolated from EDL(Extensor digitorum longus) of indicated mice and then cultured overnight. Subsequently, the cells were incubated with APC-labeled anti-FcγR1, Alexa Fluor 555-labeled-mIgG, and Alexa Fluor 488-labeled-anti-Laminin on ice overnight(**E**). The fluorescence intensity of each myofiber was determined. Data are either representative (**F**) or summary (**G, H**) of three independent experiments, and each dot represents one myofiber. Scale bars represent 100 μm.

Error bars stand for the standard deviation of the mean. * p<0.05, ** p<0.01, *** p<0.001 in two-tailed Student’s *t* test (**A**, **B, D, G, H**).

**Fig. S9. Genetic ablation of FcγR1 mitigates muscular dystrophy in *Dmd*^E4*^ mice**

(A). Ablation of FcγR1 alleviates muscular dystrophy. 6-week-old *Fcer1g^-/-^ Dmd*^E4*^ mice and *Dmd*^E4*^ mice were analyzed for remaining grip strength was determined after ten repeated tests. Data are a summary of five mice in each group, and each dot represents one individual mouse.

(B). Ablation of FcγR1 diminished tissue inflammation in *Dmd*^E4*^ mice. T.A. muscles from six to seven-week-old *Fcer1g^-/-^ Dmd*^E4*^ mice and age-matched *Dmd*^E4*^ mice were analyzed by H&E staining. Arrows indicate areas with inflammatory cell infiltration. Data are representative of three independent experiments. Scale bars represent 100 μm.

(C). Myofibers with centralized nuclei were determined in T.A. muscles. Data are summary of 5 mice per group.

(D). As in **Figure 3J**, seven-month-old *Fcer1g^-/-^ Dmd*^E4*^ mice and *Dmd*^E4*^ mice were analyzed for fibrosis of various muscle tissues with Masson’s staining. Data are a summary of at least five mice in each group, and each dot represents one individual mouse.Error bars stand for the standard deviation of the mean. * p<0.05, ** p<0.01, *** p<0.001 in two-tailed Student’s *t* test (**A**, **C, D**).

**Fig. S10. Depleting B cells using anti-CD20 antibody alleviates muscular dystrophy in DMD mice.**

**(A)** Anti-CD20 treatment in neonatal *Dmd*^E4*^ mice led to transient depletion of immunoglobulin M (IgM). One-week old *Dmd*^E4*^ mice were treated with anti-CD20 or control IgG as depicted in **Figure 5A**. Serum IgM levels were determined using enzyme-linked immunosorbent assay (ELISA). The shaded area indicates the period during which the anti-CD20 treatment was administered. Data are summarized from five to six mice in each group.

(B). Anti-CD20 treatment in neonatal *Dmd*^E4*^ mice led to hypogammaglobulinemia. As in (**A**), serum IgG levels were determined using ELISA. Data are summarized from five to six mice in each group.

(**C**) Sustained improvement of skeletal muscle functions following neonatal anti-CD20 treatment. To assess muscle strength, 10 grip tests of combined forelimb and hindlimb strength were conducted every 2-4 weeks, with a short interval (10 seconds) between each test. The remaining strength in the last cycle was calculated by normalizing to the maximal grip strength. Data are summarized from five mice in each group.

(D) Eight weeks following the anti-CD20 treatment, levels of IgG and IgM in the T.A. muscle were analyzed using immunoblotting. Data are summarized from three to six mice in each group.

(E) Anti-CD20 treatment rescued the *Dmd*^E4*^ mice from kyphosis. Kyphosis indexes (KI) were determined using μCT. Data are a summary of six mice in each group.

(F) H&E staining of R.F. and diaphragm from *Dmd*^E4*^ mice treated with anti-CD20 or control mouse IgG. Arrows indicate the area with intense inflammatory cell infiltration. Data are representative of four mice in each group. Scale bars represent 100 μm.

(G) Fibrosis was determined using Masson’s staining of the diaphragm and R.F. from *Dmd*^E4*^ mice treated with anti-CD20 or control mouse IgG as shown in (**A**). Data are representative of four to five mice in each group. Scale bars represent 100 μm.

(H) As in (**G**), fibrotic tissues were quantified based on Masson’s staining in the R.F. and Diaphragm. Data are summarized from four or five mice in each group.

Error bars stand for the standard deviation of the mean. * p<0.05, ** p<0.01, *** p<0.001, **** p<0.0001 in two-tailed Student’s *t* test (**A**, **B**, **C, E, H**).

**Fig. S11. Anti-CD20 treatment alleviates muscular dystrophy in adult *Dmd*^E4*^ and *mdx* mice.**

(A). Schematic of using anti-CD20 antibody to deplete B cells in 8-week-old *Dmd*^E4*^ mice. 8-week-old *Dmd*^E4*^ mice were injected with anti-mouse CD20 (2 mg/kg) or control mouse IgG once a week for eight weeks.

(B). Levels of circulating B cells in PBMC were determined by flow cytometry. Data are summarized from 5 to 6 mice in each group.

(C). Serum CK levels were determined prior to the treatment (week 0), and every 2-4 weeks thereafter. Gray areas indicate the time that the mice received the antibody treatment. Each dot represents one individual mouse (five to six mice in each experiment). Note that CK level reduction was sustained for at least 3 months after the last anti-CD20 treatment.

(D). As in **Figure 5D**, anti-CD20 treatment prevented contraction-induced fatigue in adult *Dmd^E4*^*mice. Two weeks after the last injection, 10 grip tests of combined forelimb and hindlimb strength were conducted with a short interval (10 seconds) between each test. The remaining strength in the last cycle was calculated by normalizing to the maximal grip strength. Data are summarized from 5 mice in each group. Each dot represents one individual mouse. (**E-G**). Anti-CD20 treatment alleviates muscular dystrophy in neonatal *mdx* mice.

(E). Schematic of using anti-CD20 antibody to deplete B cells in 1-week-old *mdx* mice. 1-week-old *mdx* mice were injected with anti-mouse CD20 (2 mg/kg) or control mouse IgG once a week for eight weeks.

(F). Levels of serum CK were determined in the indicated time. Gray area shows the time window that the mice received the antibody. Data are summarized from five mice in each group. Each dot represents one individual mouse.

(G). Eight weeks after the last treatment, 10 grip tests of combined forelimb and hindlimb strength were conducted with a short interval (10 seconds) between each test. The remaining strength in the last cycle was calculated by normalizing to the maximal grip strength. Data are summarized from five mice in each group.

Error bars stand for the standard deviation of the mean. * p<0.05, ** p<0.01, *** p<0.001, **** p<0.0001, ns, not statistically significant in two-tailed Student’s *t* test (**B, C, F**) or two-way ANOVA test (**D, G**).

**Fig. S12. Depleting B cells promotes a shift from proinflammatory into immunosuppressive milieu in dystrophic muscles**

(**A-C**). Anti-CD20 treatment shifted the proinflammatory to immunosuppressive macrophages in skeletal muscles of *Dmd* ^E4*^ mice. In 16–18-week-old *Dmd* ^E4*^ mice receiving either control IgG or anti-CD20 treatment during neonatal stages, leukocytes were isolated from skeletal muscles and M1/M2 macrophages were determined by flow cytometry. (**A**) representative F4/80 and CD11b staining, which was pregated on CD45+ cells. (**B**), representative Ly6C and CD206 staining of pregated macrophages. M1-like proinflammatory macrophages were characterized by CD206^low^ Ly6C^hi^ and immunosuppressive macrophages (M2-like) were CD206^hi^Ly6C^low^. (**C**). Summary of numbers of infiltrating leukocytes, total macrophages, percentage of M1/M2 macrophages in the skeletal muscles. Each dot represents one individual mouse. Error bars stand for the standard deviation of the mean. * p<0.05, ** p<0.01, *** p<0.001 in two-tailed Student’s *t* test.

(**D, E**). scRNA seq analysis of immune cell composition in the skeletal muscles of *Dmd*^E4*^ mice. Neonatal *Dmd* ^E4*^ mice were treated with anti-CD20 or control IgG as illustrated in **Figure 5A**. 12 weeks after the last injection, total CD45^+^ cells in skeletal muscles were sorted by flow cytometry and analyzed by scRNA seq.

(**D**). UMAP analysis showing all cells isolated from the anti-CD20 (n=12227) and IgG-(n=16655) treated mice, including a total of 11 clusters. Four clusters (0,1,2,3) belong to the mono/macrophages (Mon/Mac). MSC, mesenchymal stromal cells; cDC1, conventional type 1 Dendritic Cells.

(**E**). Dot plot shows the cell type-specific markers for clusters identified in (**D**). Top differentially expressed genes that were upregulated in each cluster were plotted.

(**F**). Lack of a clear correlation between M1/M2 signatures in scRNA-seq identified clusters. Expression of M1 and M2 macrophage markers in M1-like macrophages (cluster 0), C1q-M2 like macrophages (cluster 1), and proliferating macrophages (cluster 3) from scRNA seq results.

(**G**). Violin plots comparing gene expression of skeletal muscle macrophages between control IgG and anti-CD20-treated *Dmd*^E4*^ mice across five identified clusters. Genes associated with GO0002683 (negative regulation of immune system process) and GO0006954 (inflammatory response) were shown. Color-coded numbers at the bottom indicate clusters identified in **Figure 5H**.

(**H-J**) The interstitial fluid collected from the skeletal muscles of the indicated mice was subjected to Olink analysis to identify differentially accumulated inflammatory factors (p<0.01 and LFC>0.25). The data are summarized from four wild type mice, six *Dmd*^E4*^ mice treated with control IgG, and six *Dmd*^E4*^ mice treated with anti-CD20 (**H**). (**I,J**).Ccl2 and IL-10 levels were determined using Olink as in (**H**).

