## supplemental file for "Elimination of intramuscular immunoglobin accumulation alleviates Duchenne Muscular Dystrophy"

#### Materials and Methods

Further information and requests for resources and reagents should be directed to and will be fulfilled by the corresponding author Xing Chang. All unique/stable reagents generated in this study are available with a completed Materials Transfer Agreement.

##### *Mice*

C57BL/10-mdx mice, uMT mice were obtained from the Jackson Laboratory. C57BL/10- *Dmd* $^{E4*}$ mice were originally from National Resource Center of Model Mice (Nanjing, China). C57BL/10-*mdx* mice and C57BL/10-*Dmd* $^{E4*}$ mice were backcrossed to C57BL/6 background for at least ten generations. ROSA26-LoxP-STOP-LoxP-tdTomato (Ai9) mice and Lyz2-cre mice were kindly provided by Dr. Heping Xue and Dr. Yi-chuan Xiao (Shanghai Institute of Nutrition and Health, Chinese Academy of Sciences) respectively. Fc receptor common γ-chain-deficient mice (Fcer1g$^{-/-}$)(*66*) were kindly provided by Dr. Jeffrey Ravetch (The Rockefeller University). Because *Dmd* gene is located on the X chromosome, only male mice were used in the study. All animal studies were performed in compliance with guidelines for the care and use of laboratory animals and were approved by Westlake University's Institutional Biomedical Research Ethics Committee.

##### *NHP*

The Rhesus monkeys utilized in this study were accommodated in a controlled environment, with a temperature of 22 ± 1 °C and humidity levels maintained at 50% ± 5% RH. They were exposed to a consistent 12-hour light/dark cycle with lights being switched on at 08:00 a.m. Their diet consisted of a commercial monkey blend, fed twice daily, supplemented with fruits and vegetables served once per day. The animal materials used and experimental procedures performed were approved by the Institutional Animal Care and Use Committee of the Yunnan Key Laboratory of Primate Biomedical Research (IACUC number: LPBR202103005).

##### *Muscle force measurement of Dmd mice*

Muscle force measurement was performed as previously described[(12)] using a grip strength meter (SA417, Jiangsu SANS Biological Technology, China), and only peak muscle force was recorded in each cycle.

##### *Micro-CT*

Micro-CT measurement was performed as previously described[(12)]. Mice were anesthetized, and the kyphosis was evaluated by μ-computed tomography (Skyscan 1276, Bruker). The KI was calculated as previously described[(67)].

##### *Cardiac function analysis*

Echocardiography was performed using Vevo3100 small animal ultrasound imaging system (Visual Sonics) equipped with a 70-MHz imaging transducer. Anesthetized mice were placed on the heated platform (34℃), and touched with ultrasound probe. Two-dimensional B-mode imaging was used to capture the long-axis projections and LVEF and LVFS were measured in M-Mode. Mean values for each echocardiogram parameter were collected from five consecutive cardiac cycles.

***IgG/M transfer***

Purified mouse IgG (Absin abs20038) or purified mouse IgM (CELLWAYLAB C040311) were administered to 1-week-old *Dmd* $^{E4*}$ uMT mice (100μg/e.a.) twice a week via intraperitoneal injection for two weeks. Subsequently, the mice were treated with purified mouse IgG or IgM (300μg/e.a.) twice a week for another four weeks.

***Myofiber isolation, culture, and immunofluorescence staining***

The isolation of myofibers was performed as previously described(*68*). Briefly, EDL (extensor digitorum longus) muscles were dissociated in a digestion buffer containing 2 mg/mL of collagenase type I (LS004196, Worthington) in serum-free DMEM media (Hyclone), with constant shaking at 37°C for 90-150 minutes. To stop digestion, the muscles were transferred to a prewarmed Petri dish containing 1 mL of DMEM using a large bore glass pipette. The muscles were then dissociated with the same pipette until the desired number of myofibers were floating freely in the solution. Subsequently, a small-bore glass pipette, coated with HS (horse serum), was used to transfer approximately 50-100 myofibers onto Matrigel (Corning, 354277) coated dishes. These dishes were pre-filled with myofiber culture medium, which consisted of DMEM supplemented with 20% FBS, 10% HS, 1% Chicken Embryo extract（Absin，abs80002）, and 1% Penicillin/Streptomycin. The myofibers were then incubated at 37°C in a 5% CO2 environment.

Adherent myocytes were fixed using a 4% solution of paraformaldehyde (PFA)， permeabilized in 0.4% Triton X-100 for 15mins, and blocked in 4% BSA. Primary antibodies used for immunofluorescence staining were as follows: APC anti-mouse CD64 (FcγRI) (161005, Biolegend), anti-Laminin (L9393, Sigma), Purified mouse IgG (Absin abs20038) and Purified mouse IgM (CELLWAYLAB C040311). Images were obtained with Olympus FV3000 Confocal Microscope and Nikon Motorized Fluorescence Microscope. The Integrated density of each channel was measured with Image J software based on the density of pixels.

***Givinostat treatment***

Myofibers derived from 8-week-old *Dmd* $^{E4*}$ mice were plated in 24-well plates at a density of 50 to 100 cells per well. The cultures were then treated with Givinostat (MedChemExpress) with indicated concentrations.

***AAV9 production and injection***

TAM-CBE (AIDx*-CO-nsaCas9(KKH)-Ugi AAV9) and sgRNA (5' SS-E4-Dmd sgRNA) vectors were packaged with AAV9 capsid as described (*12*) by PackGene Biotech (Guangzhou, China). ShRNAs targeting FcγR1 (shFcgr1-1, 5′-gatCCGGTCACGGTGAAAGAGCTGTTTACTCGAGTAAACAGCTCTTTCACCGTGATTTTTG-3′;shFcgr1-2, 5′-gatCCGGGCAAATTCCTTTCAGCAAGTTCTCGAGAACTTGCTGAAAGGAATTTGCTTTTTG-3′) were cloned into a scAAV vector under the U6 promoter followed by a CAG promoter-driven EGFP cassette. The vector were packaged with AAV9 capsid by PackGene Biotech (Guangzhou, China). The tibialis anterior (T.A.) muscle of 4-week-old *Dmd* $^{E4*}$ mice was injected with scAAV-shRNAs particles against FcγR1 (5x10^11 vg per T.A.), and FcγRI expression and IgG accumulation in treated T.A. were assessed and were compared to untreated T.A. on the opposite side in the same mice.

***OLINK analysis***

Muscle tissue interstitial fluid (TIF) was obtained via serial centrifugation as described (*69*). Briefly, the tibialis anterior (TA) muscles were isolated and positioned atop a 40-micron strainer. This strainer was then secured within a 50 mL Falcon tube. The centrifugation process was conducted in two stages: an initial spin at 50×g for 5 minutes at 4°C to remove any residual surface liquid, followed by a secondary spin at 600×g for 10 minutes at 4°C to collect the TIF. The collected fluid was subsequently prepared for further processing and analysis.

Following the collection of muscle tissue interstitial fluid (TIF), protein levels were measured utilizing the Olink Target 96 Mouse Exploratory Panel (Olink Proteomics AB, Uppsala, Sweden), in accordance with the manufacturer's protocol. The data were normalized using both an internal extension control and an inter-plate control, which adjusts for any intra- and inter-run variations that may occur. The outcome of the assay is expressed in Normalized Protein eXpression (NPX) values, which are arbitrary units on a log2 scale, with higher values indicating increased protein expression.

***B cell depletion with anti-mouse CD20***

Anti-mouse CD20 (BE0356, Bio X Cell, Clone: MB20-11) and mouse IgG control (BE0366, Bio X Cell, Clone: DV5-1) were administered to 1-week-old or 8-week-old *Dmd* $^{E4*}$ mice once a week via intraperitoneal injection (2mg/kg) for 8 weeks.

***Anti-CD20 (Rituximab) treatment in DMD monkeys***

Rituximab (Meilunbio)(20mg/kg) was diluted in 50 ml of 0.9% normal saline and administered via an intravenous drip to DMD monkeys once a week for 4 weeks. WT monkeys and age matched DMD monkeys that received saline were included as controls.

***Tests for monkey exercise capacities and motor functions***

***Locomotion***

The experimental monkey was transferred to a clean, transparent standard video cage in a separate room to avoid interference from other monkeys. The monkey can move freely in the cage and have free access to drinking water. No one was in the room after starting recording to avoid interference. The monkey activity was recorded continuously with a Kinect 2.0 camera for 60 min and analyzed by Primate Scan 1.0 software (Clever Sys Inc, VA).

***Lie-to-stand assay***

This experiment was adapted from Govers' sign, a typical feature of DMD patients. Monkey's hands were tied in front of or behind its body and the monkey were then placed in a prone position. Each DMD or WT monkey was recorded at least three times.

***Magnetic Resonance Imaging***

Monkeys were first anesthetized and positioned in supine in the bore of a Siemens Magnetom Prisma 3T MRI Machine. Multi-sequence images of lower limbs were collected. The results were subsequently analyzed using a DICOM Viewer to identify any indications of fatty accumulation, inflammation, and edema.

***Immunoblotting***

To detect protein expression, tissues were lysed with RIPA buffer (50mM Tris-HCl pH7.4, 150mM NaCl, 1mM EDTA, 1% Triton-X100, 1% sodium deoxycholate, 0.1% sodium dodecyl sulphate (SDS) supplemented with complete protease inhibitors and 1mM DTT). Proteins were separated on 6 or 10% SDS-PAGE, and transferred onto polyvinylidene fluoride (PVDF) sheets. Immunoglobins in the skeletal muscles were detected using horseradish peroxidase (HRP)-labeled antibodies: Goat anti-mouse IgG (SouthernBiotech, 1030-05) and Goat anti-mouse IgM (SouthernBiotech, 1020-05). Other primary antibodies used for immunoblotting were as follows: Goat Anti-Mouse IgG1, Human ads-HRP (1070-05, SouthernBiotech), Goat Anti-Mouse IgM, Human ads-HRP(1020-05, SouthernBiotech)，Goat Anti-Mouse IgG2b, Human ads-HRP (1090-05, SouthernBiotech)，Goat Anti-Mouse IgG2c, Human ads-HRP(1079-05, SouthernBiotech)，Goat Anti-Mouse IgG3, Human ads-HRP(1100-05, SouthernBiotech)，anti-Vinculin (ab129002, Abcam), and anti-GAPDH (sc-48166, Santa Cruz).

***Immunofluorescence staining***

Anesthetized mice were first perfused with PBS, and hearts were rinsed with cold PBS containing 2M KCl. These fresh tissues were immediately embedded in O.C.T. Compound (4583, SAKURA) and kept frozen in -80℃ freezer. The embedded issues were cut at 10 μm intervals and were collected on frost-coated slides. For Immunofluorescence staining, the slides were washed with PBS, fixed with 4% paraformaldehyde for 10mins, permeabilized in 0.4% Triton X-100 for 15mins, and blocked in 4% BSA. Primary antibodies used for immunofluorescence staining were as follows: Goat anti-Mouse IgG (H+L) (Alexa Fluor™ 555) (A-21424, Invitrogen), Goat anti-Mouse IgM mu chain (Alexa Fluor® 488) (ab150121, Abcam), anti-Laminin (L9393, Sigma), Donkey anti-Rabbit IgG (H+L) (Alexa Fluor™ 647) (A-31573, Invitrogen), Donkey anti-Rabbit IgG (H+L) (Alexa Fluor™ 488) (A-21206, Invitrogen), APC anti-mouse CD64 (FcγRI) (161005, Biolegend), Anti-Dystrophin(PA532388, Invitrogen), Alexa Fluor® 488-AffiniPure Alpaca Anti-Human IgG (H+L) (609-545-213, Jackson ImmunoResearch Laboratories), APC anti-human IgM (314510, Biolegend). The nuclei were stained with DAPI (abs47047616, Absin). Images were obtained with Olympus FV3000 Confocal Microscope and Nikon Motorized Fluorescence Microscope. And the Integrated density of each channel was measured with Image J software based on the density of pixels.

***Serum Creatine Kinase (CK) measurement***

Creatine Kinase levels in serum were assessed using a Creatine Kinase-SL ASSAY (326-10, SEKURE CHEMISTRY) according to the manufacturer's instructions. Blood was collected via the submandibular vein method. Serum was isolated and kept frozen in -80℃ freezer.

***Immunoprecipitation and Mass spectrometry analysis***

Skeletal muscles were lysed with RIPA buffer (50mM HEPES pH7.5, 150mM NaCl, 10% glycerol, 0.2% Triton-X100, supplemented with complete protease inhibitors), and the supernatants were collected. Protein A/G beads were added to each sample for 3hr incubation at 4°C and samples were washed 5 times in wash buffer (50mM HEPES pH7.5, 150mM NaCl, 0.1% Triton-X100, supplemented with complete protease inhibitors). Immunoprecipitates were eluted by boiling for 10 minutes in diluted loading buffer (P0015F, Beyotime), and were used for mass spectrometry detection. The MS data have been deposited in the ProteomeXchange proteomics Consortium (http://proteomecentral.proteomexchange.org) via the PRIDE partner repository with the following accession number: PXD052136 (Reviewer Username:, Password: YEvJngHY).

***Flow cytometry analysis of peripheral blood mononuclear cells (PBMC)***

Peripheral blood was collected via submandibular vein method from the anti-CD20-treated *Dmd* $^{E4*}$ mice and the control antibody-treated mice, and red blood cells were removed by RBC lysis

(C3702, Beyotime). For surface staining, cells were blocked with an antibody against CD16/32 (2.4G2, BD Pharmingen), and indicated with LIVE/DEAD (L34994, Invitrogen) and indicated antibodies for 30min at 4°C. The cells were then washed with staining buffer (1% FBS in PBS) and were analyzed with a CytoFLEX flow cytometer and CytExpert software according to the manufacturer's instructions. The following antibodies were used in this study. Anti-mouse CD45-APC/Cyanine7 (30-F11, Biolegend, 103115), anti-mouse CD19- FITC (1D3/CD19, Biolegend, 152403). Anti-human CD20- FITC (B-LY1, Santa Cruz, sc-19990) was used to detect B cells in PBMC of monkeys.

***Isolation and analysis of muscle mononuclear cells***

The isolation of mononuclear cells in the muscle was performed as previously described[70]. Briefly, mice were first perfused with PBS and lymph nodes were removed before dissecting skeletal muscles (Tibialis Anterior, Rectus Femoris and Diaphragm) to exclude the contamination of non-muscle-residing lymphocytes. Minced muscle tissues were dissociated in digestion buffer containing 1 mg/mL collagenase type II (LS004177, Worthington), 20 μg/mL DNase I (11284932001, Roche), and 10% FBS (Hyclone) in serum-free DMEM media (Hyclone) with constant stirring at 37 °C for 30 min. Digested tissues were sequentially filtered through a 100μm filter-basket. Mononuclear cells were then collected at the interface of a 40%–70% Percoll gradient (17089109, cytiva), and analyzed by Spectra Analyzer-Aurora (Cytek).

The following FACS antibodies were used in the study: anti-Mouse CD16/32 (2.4G2, BD Pharmingen™, 553142), anti-Mouse CD45-BUV395 (30-F11, BD Pharmingen™, 564279), anti-mouse NK-1.1-Brilliant Violet 650™ (PK136, Biolegend, 108735), anti-mouse CD170 (Siglec-F)-PE (S17007L, Biolegend, 155505), anti-mouse Ly-6C-FITC (HK1.4, Biolegend, 128005), anti-mouse F4/80-PE/Cyanine7 (BM8, Biolegend, 123113), anti-mouse/human CD11b-PerCP/Cyanine5.5 (M1/70, Biolegend, 101227), anti-mouse Ly-6G-Pacific Blue (1A8, Biolegend, 127611), anti-mouse CD206-Alexa Fluor® 647 (1A8, BD Pharmingen™, 565250), anti-mouse CD3 -eFluor™ 660 (17A2, Invitrogen, 50003282), anti-Mouse TCR β-BV605 (H57-597, BD Pharmingen™, 562840), anti-Mouse γδ T-Cell Receptor -BV605 (GL3, BD Pharmingen™, 561997), anti-mouse CD4-Brilliant Violet 510™ (GK1.5, Biolegend, 100449), anti-mouse CD8a-PE/Cy7 (53-6.7, BD Pharmingen™, 561097), anti-mouse CD19-PerCP/Cyanine5.5 (1D3/CD19, Biolegend, 152405), anti-mouse/human B220-FITC (RA3-6B2, Biolegend, 103205), Foxp3-eFluor™ 450 (FJK-16s, Invitrogen, 48577382).

***ELISA***

ELISA plate (F605031-0001, BBI) was coated with Goat anti-mouse Ig (1010-01, SouthernBiotech) by incubating overnight at 4°C. Then, the plate was blocked in 3% BSA. Next, samples were added to the plate, which was followed by adding the detection antibody (Anti-mouse IgG-HRP (0107-01, SouthernBiotech), Goat Anti-Mouse IgM-HRP (1020-05, SouthernBiotech)). The chromogenic reaction which converted the substrate (TMB) into a colored product was measured using a plate reader.

***Histology analysis***

Skeletal muscles and hearts were washed in PBS, fixed in 10% neutral formalin, embedded into paraffin after dehydration and sectioned (8μm) for further analysis. Hematoxylin and Eosin staining was conducted following the manufacturer's protocol (G11220, Solarbio). Briefly, dewaxing of the tissue sections was performed using xylene, repeated twice for a duration of 5 minutes each. Following dewaxing, rehydration was carried out sequentially with 100% ethanol, 90% ethanol, and 70% ethanol for 3 minutes per solution. Staining with hematoxylin was conducted for 15 minutes, after which the sections were immersed in a 1% hydrochloric acid (HCl) solution for 1 second. This was followed by staining with a 1% eosin solution for 2 minutes. Finally, dehydration of the sections was achieved using a graded series of ethanol solutions: 70% ethanol, 90% ethanol, and 100% ethanol. Finally, the sections were soaked in xylene and mounted with a coverslip using mounting medium. Stained samples were analysed with Motorized Fluorescence Microscope (Ni-E, Nikon).

Masson's staining was conducted following the manufacturer's protocol (G2340, Solarbio) to determine the degree of muscle fibrosis, and Weigert's hematoxylin, Biebrich scarlet-acid fuchsin solution, and aniline blue were utilized for the detection of collagen fibers within tissues. Briefly, Weigert's hematoxylin and the Biebrich scarlet-acid fuchsin solution were applied for staining, each for a duration of 10 minutes. This process results in the muscle cells appearing red. Subsequently, aniline blue was used for staining for 5 minutes. Fibrotic tissues were stained blue, and the fibrosis ratio was measured with Image J software based on the density of pixels.

***Single-molecule RNA in-situ hybridization***

The expression of FcγR1 mRNA on myofibers was evaluated using an RNAscope Assay (323100, Advanced Cell Diagnostics (ACD)). Frozen tissue samples were prepared according to the manufacturer's protocol and pretreated with RNAscope® Protease IV (322381, ACD). An RNAscope probe targeting the specific sequences of FcγR1 was designed (Mm-FcγR1Lot: 23111B). The probe signal was amplified by AMP1, AMP2, and AMP3 (323110, ACD). RNAscope assays were performed using the Opal™ 570 detection reagent (FP1488001KT, ACD) per the manufacturer's protocol. The nucleus and cell membrane were labeled with DAPI and Laminin (Alexa Fluor™ 488).

***Preparation of Single Cell 3' Gene Expression dual index libraries***

CD45-positive cells in the skeletal muscles of *Dmd* $^{E4*}$ mice treated with either anti-CD20 or control IgG were sorted with flow cytometry, and Gel Beads in emulsion (GEMs) were generated by combining barcoded Single Cell 3' v3.1 Gel Beads, a Master Mix containing cells, and Partitioning Oil onto Chromium Next GEM Chip G. Single Cell 3' Gene Expression dual index libraries was prepared using Chromium Next GEM Chip G Single Cell Kit (10xGenomics, PN-1000120) according to manufacturer's instruction. 10X Genomic libraries were sequenced using Illumina Nova Seq 6000 platform.

***Single-cell raw data processing and analysis***

Illumina basecall files (*.bcl) were converted to fastqs using the Cell Ranger v6.1.2 pipeline with recommended parameters. Each library was aligned to an indexed mm10 genome using Cell Ranger Count. This output matrix was then imported into the Seurat (version 4.0.2) R toolkit for quality control and downstream analysis. After removing the double cells with scrublet (v. 0.2.3)(*71*), the Seurat pipeline (version 4.3.01) was applied to the aligned cell matrix using R (version 4.2.1) to identify cell clusters. Quality control filtering removed genes that were not expressed (>0) in at least three cells and cells with less than 300 genes; red blood cell contamination was subsequently removed. Seurat's default clustering was performed and followed with marker gene detection to elucidate gene expression signatures corresponding to the resultant clusters.

***Reference-based mapping of macrophages***

Anchoring and integration were first performed for single-cell datasets of immune cells isolated from control IgG or anti-CD20 treated *Dmd* $^{E4*}$ mice. Seurat (version 4.3.01, R version 4.2.1) was used for anchoring and integration (*72*). Briefly, merged Seurat objects were normalized, and highly variable genes (features) and scaling were performed with SCTransform(*73*). The top 3000 highly variable features were selected and used for anchoring. Integration anchors (50 dimensions) were computed and used for integration. For neighbor and cluster identification, the integrated object was scaled, and significant PCs were identified via statistical and heuristic testing as recommended in Seurat. Clustered cells were visualized using UMAP. Before anchoring and integration, macrophage identities from both datasets were renamed to match macrophage identities from the current study. All other previously assigned cell identities described macrophage were kept in the final clustering. Next, all identified macrophages were used for second-round clustering with the same procedure for further analysis.

***Identification of differentially expressed genes (DEG)***

DEGs were determined based on the Wilcoxon rank-sum test that was implemented in the function FindMarkers and FindAllMarkers in the Seurat package. Unless noted otherwise, genes with $P < 0.05$, absolute log2 fold change >0.25, and minimum fraction >0.1 were selected as DEGs.

***Enrichment analysis for signature genes***

Metascape (https://metascape.org) was used to perform the enrichment analysis for multiple DEGs lists(*74*).

***Statistics***

Statistical tests were performed using two-tailed Student's t tests or 2-way ANOVA tests in GraphPad Prism version 8.0 unless specified in the manuscript. Error bars stand for the standard deviation of the mean unless specified. *, $p < 0.05$; **, $p < 0.01$; ***, $p < 0.001$; ****, $p < 0.0001$ in two-tailed Student's t tests or 2-way ANOVA tests.

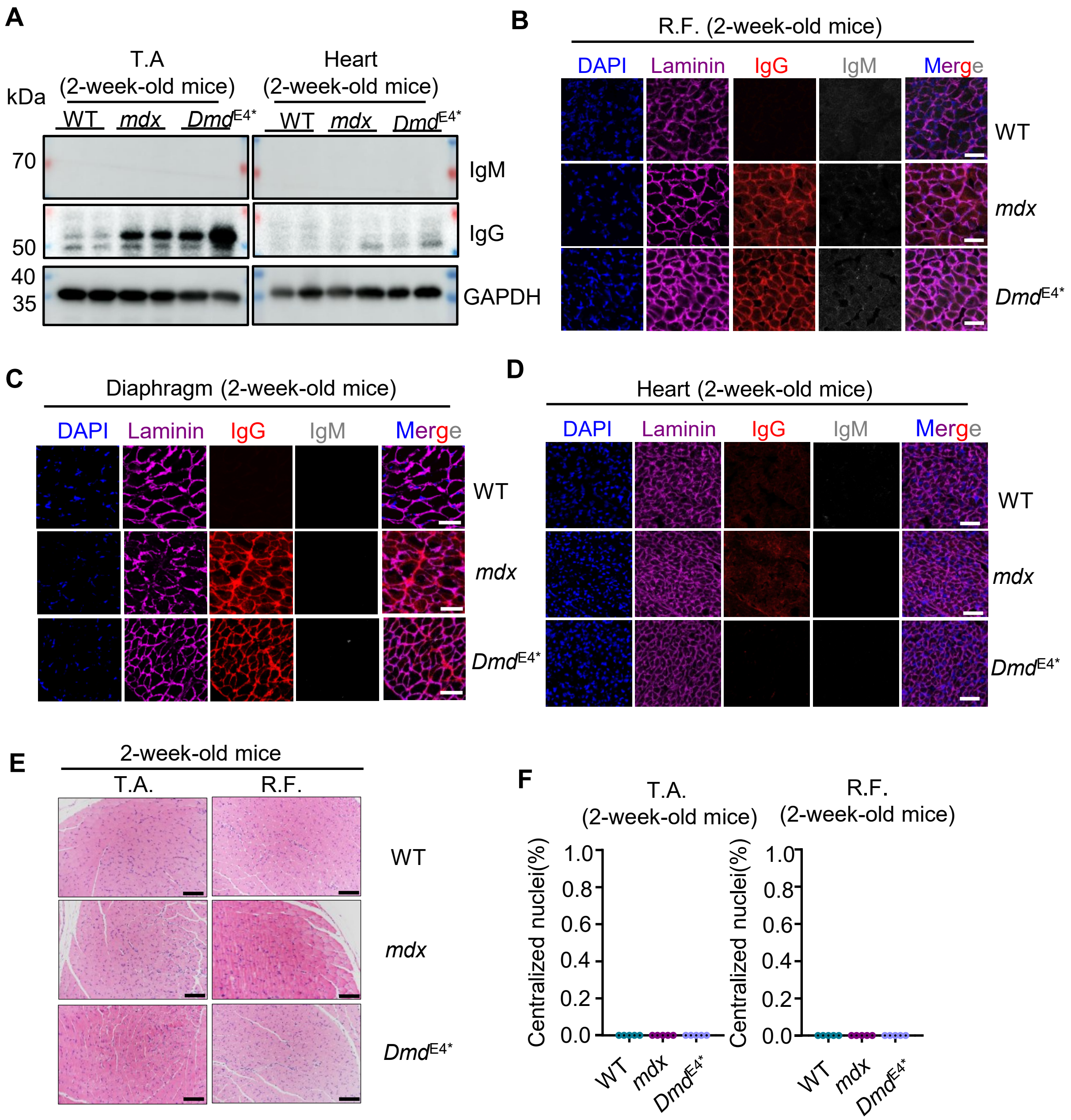
A
T.A (2-week-old mice)
Heart (2-week-old mice)
kDa
WT mdx DmdE4*
70
50
40
35
IgM
IgG
GAPDH
B
R.F. (2-week-old mice)
DAPI Laminin IgG IgM Merge
WT
mdx
DmdE4*
C
Diaphragm (2-week-old mice)
D
Heart (2-week-old mice)
E
2-week-old mice
T.A.
R.F.
F
T.A. (2-week-old mice)
R.F. (2-week-old mice)
Centralized nuclei(%)
1.0
0.8
0.6
0.4
0.2
0.0

**Figure S1**

**Fig. S1. Accumulation of IgG in various skeletal muscles of 2-week-old *Dmd*$^{E4*}$ and *mdx* mice**

(**A**). Tissue lysate from 2-week-old *mdx*, *Dmd*$^{E4*}$, and WT mice was analyzed by immunoblotting for IgM and IgG. GAPDH was included as loading controls. Data are representative of six mice in each group.

(**B-D**). Tissue sections of rectus femoris (R.F.) (**B**), diaphragm (**C**), and heart (**D**) (10 μm thick) were stained with Alexa Fluor 555-labeled anti-mouse IgG, Alexa Fluor 647-labelled anti-Laminin, Alexa Fluor 488-labeled anti-mouse IgM, and DAPI. Data are representative of five mice in each group. Scale bars represent 50 μm.

(**E-F**). There were no detectable inflammatory cell infiltration and muscles with centralized nuclei in 2-week-old *mdx* and *Dmd*$^{E4*}$ mice. (**E**). Tissue sections of T.A. and R.F. muscles were examined with H&E staining. Scale bars represent 100 μm. (**F**). Summary of Myofibers with centralized nuclei in T.A. and R.F. muscles. Data are summarized from five mice in each group.

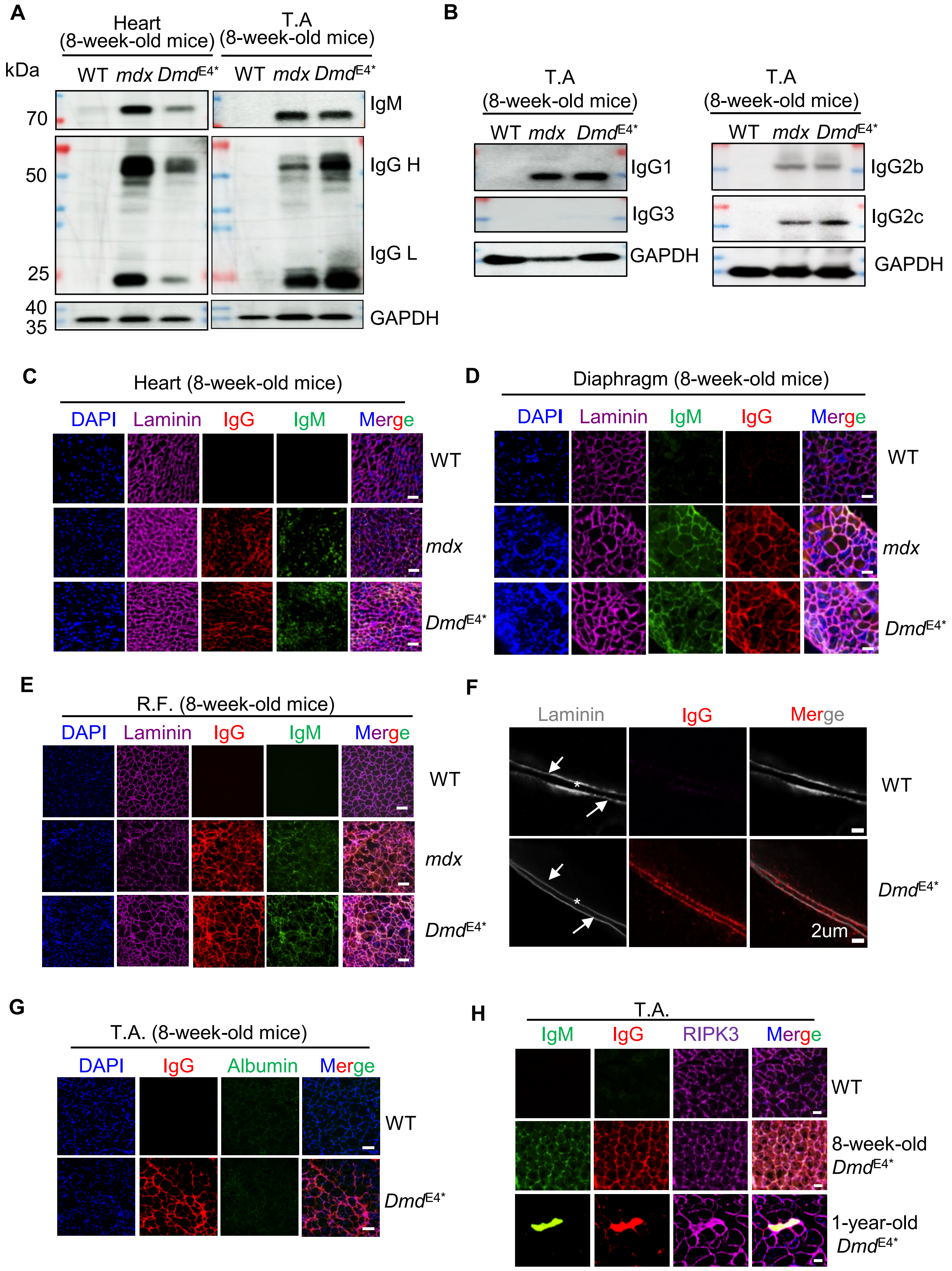
A
Heart (8-week-old mice)
T.A (8-week-old mice)
kDa
WT mdx DmdE4*
WT mdx DmdE4*
70
50
25
40
35
IgM
IgG H
IgG L
GAPDH
B
T.A (8-week-old mice)
WT mdx DmdE4*
IgG1
IgG3
GAPDH
IgG2b
IgG2c
C
Heart (8-week-old mice)
DAPI Laminin IgG IgM Merge
WT
mdx
DmdE4*
D
Diaphragm (8-week-old mice)
DAPI Laminin IgM IgG Merge
E
R.F. (8-week-old mice)
F
Laminin IgG Merge
2um
G
T.A. (8-week-old mice)
DAPI IgG Albumin Merge
H
T.A.
IgM IgG RIPK3 Merge
8-week-old DmdE4*
1-year-old DmdE4*

**Figure S2**

**Fig. S2. Accumulation of immunoglobins in skeletal and cardiac muscles of 8-week-old DMD mice**

(**A**). Immunoblot analysis of IgG accumulation. Total tissue lysate from the heart (left panel) or T.A. (right panel) was analyzed for IgM, IgG heavy chain (IgG H), or IgG light chain (IgG L). Data are representative of six mice in each group.

(**B**). Levels of IgG1, IgG2b, IgG2c, and IgG3 were analyzed in tissue lysate of T.A. muscles as in (**A**). Data are representative of four to six mice in each group. GAPDH was included as a loading control, and a molecular weight standard was marked on the left.

(**C-E**). Tissue sections of the heart (**C**), diaphragm (**D**), and R.F. (**E**) were stained with Alexa Fluor 555-labeled anti-mouse IgG, Alexa Fluor 647-labeled anti-Laminin, Alexa Fluor 488-labeled anti-mouse IgM. Data are representative of five mice in each group. Scale bars represent 50 μm.

(**F**). Super-resolution microscopy imaging of skeletal muscle tissues. The sarcolemma was stained with Laminin (gray) and indicated by arrows. The endomysial space, which represents the interstitial space between adjacent sarcolemmas, is highlighted by asterisks. Data are representative of four mice.

(**G**) Lack of albumin accumulation in the 8-week-old DMD muscles. T.A. muscles from 8-week-old $Dmd^{E4*}$ mice were analyzed for the presence of albumin and IgG by immunofluorescence staining. Data are representative of five mice in each group. Scale bars represent 100 μm.

(**H**). Most myofibers with IgG accumulation did not show RIPK3 aggregation in 8-week-old $Dmd^{E4*}$ mice. Immunofluorescence staining showed the subcellular localization of RIPK3 in the T.A. of WT mice, 8-week-old $Dmd^{E4*}$ mice, and 1-year-old $Dmd^{E4*}$ mice. Notably, only in the myofibers of 1-year-old $Dmd^{E4*}$ mice did the distribution of RIPK3 change to a more punctate and aggregated distribution, indicating the formation of necrosomes. Note, myofibers with RIPK3 punctate also had cytoplasmic IgM and IgG accumulation. Data are representative of three mice in each group. Scale bars represent 50 μm.

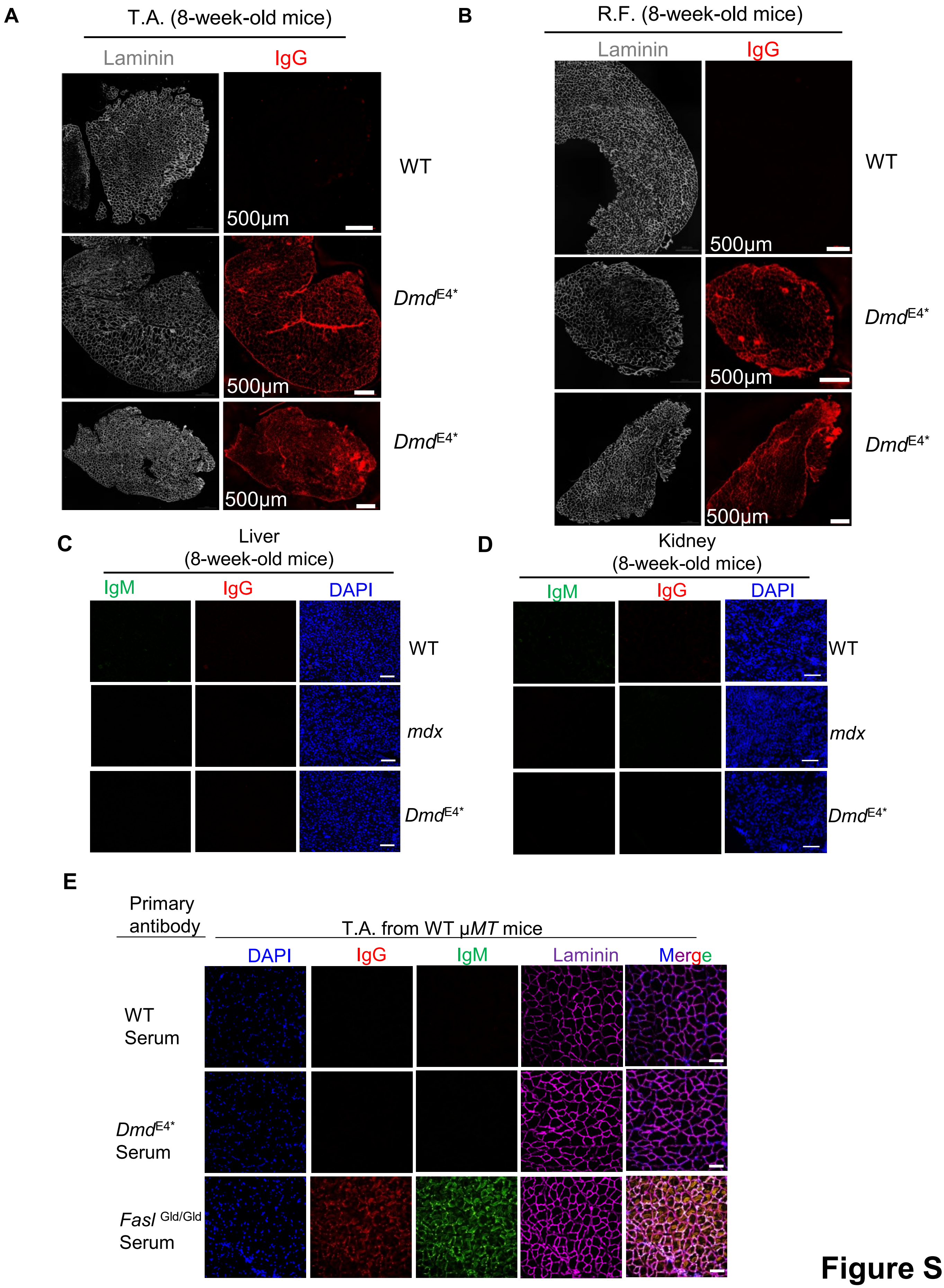
A
T.A. (8-week-old mice)
Laminin
IgG
WT
500μm
DmdE4*
500μm
DmdE4*
500μm
B
R.F. (8-week-old mice)
Laminin
IgG
WT
500μm
DmdE4*
500μm
DmdE4*
500μm
C
Liver
(8-week-old mice)
IgM
IgG
DAPI
WT
mdx
DmdE4*
D
Kidney
(8-week-old mice)
IgM
IgG
DAPI
WT
mdx
DmdE4*
E
Primary antibody
T.A. from WT μMT mice
DAPI
IgG
IgM
Laminin
Merge
WT Serum
DmdE4* Serum
Fasl Gld/Gld Serum

**Figure S3**

**Fig. S3. Characterization of antibody accumulation in the 8-week-old *Dmd* mice**

(**A,B**). Mosaic micrographs of the entire cross section of T.A. (**A**) or R.F. (**B**) from 8-week-old $Dmd^{E4*}$ mice and WT mice. Data are representative of five mice in each group.

(**C,D**). Lack of antibody accumulation in non-muscular tissues of *mdx* or $Dmd^{E4*}$ mice. Liver (**C**) and kidney (**D**) sections from 8-week-old WT, *mdx*, and $Dmd^{E4*}$ mice were analyzed for the presence of IgG and IgM by immunofluorescence staining. Data are representative of five mice in each group. Scale bars represent 100 μm.

(**E**). Serum from 2-week-old $Dmd^{E4*}$ mice or WT littermates were used as the primary antibody to stain T.A. tissue sections (10 μm thick) from WT μMT mice, which was followed by staining with Alexa Fluor 555-labeled anti-mouse IgG and Alex Fluor 488-labeled anti-mouse IgM. Data are representative of five mice in each group. Serum from lupus mice ($FasL^{gld/gld}$) was included as positive controls. Scale bars represent 100 μm.

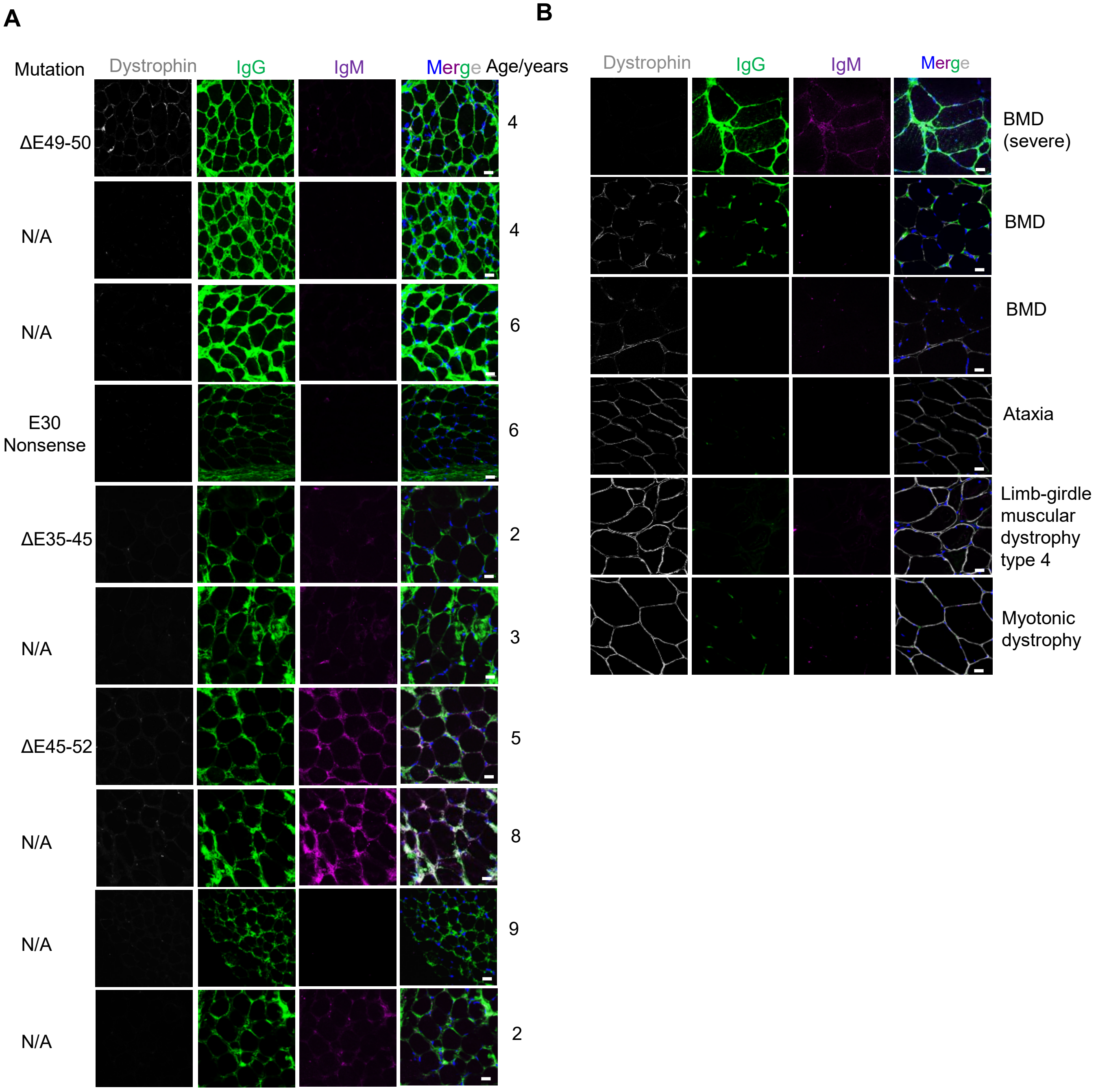

A
Mutation
Dystrophin
IgG
IgM
Merge
Age/years
ΔE49-50
4
N/A
4
N/A
6
E30 Nonsense
6
ΔE35-45
2
N/A
3
ΔE45-52
5
N/A
8
N/A
9
N/A
2
B
Dystrophin
IgG
IgM
Merge
BMD (severe)
BMD
BMD
Ataxia
Limb-girdle muscular dystrophy type 4
Myotonic dystrophy


**Figure S4**

**Fig. S4. IgG accumulation at sarcolemma of patients with Duchenne and Becker Muscular Dystrophy**

(**A**). IgG accumulation in patients with DMD. Muscle biopsies from patients with DMD at various ages were examined for the expression of dystrophin and the accumulation of IgG and IgM. The ages and DMD mutations of the patients were indicated on the right and left respectively. Tissue sections (10 µm thick) were stained with Alexa Fluor 488-labeled anti-human IgG, Alexa Fluor 594-labelled anti-Dystrophin, Alexa Fluor 647-labeled anti-human IgM, and DAPI. Each slide represents one patient. Scale bars represent 20 µm. N.A., genetic information of the patients was not available due to privacy regulations or because the patients were diagnosed before the widespread adoption of genetic testing.

(**B**). IgG accumulation in patients with BMD. Muscle biopsies from patients with BMD were analyzed for the expression of Dystrophin and the accumulation of IgG and IgM. Note a significant accumulation of IgG was observed in a patient with severe BMD with greatly reduced Dystrophin. Muscle biopsies from patients with Ataxia, Limb-girdle muscular dystrophy (type 4), and myotonic dystrophy were also included in the analysis. Tissue sections (10 µm thick) were stained with Alexa Fluor 488-labeled anti-human IgG, Alexa Fluor 594-labelled anti-Dystrophin, Alexa Fluor 647-labeled anti-human IgM, and DAPI. Scale bars represent 20 µm.

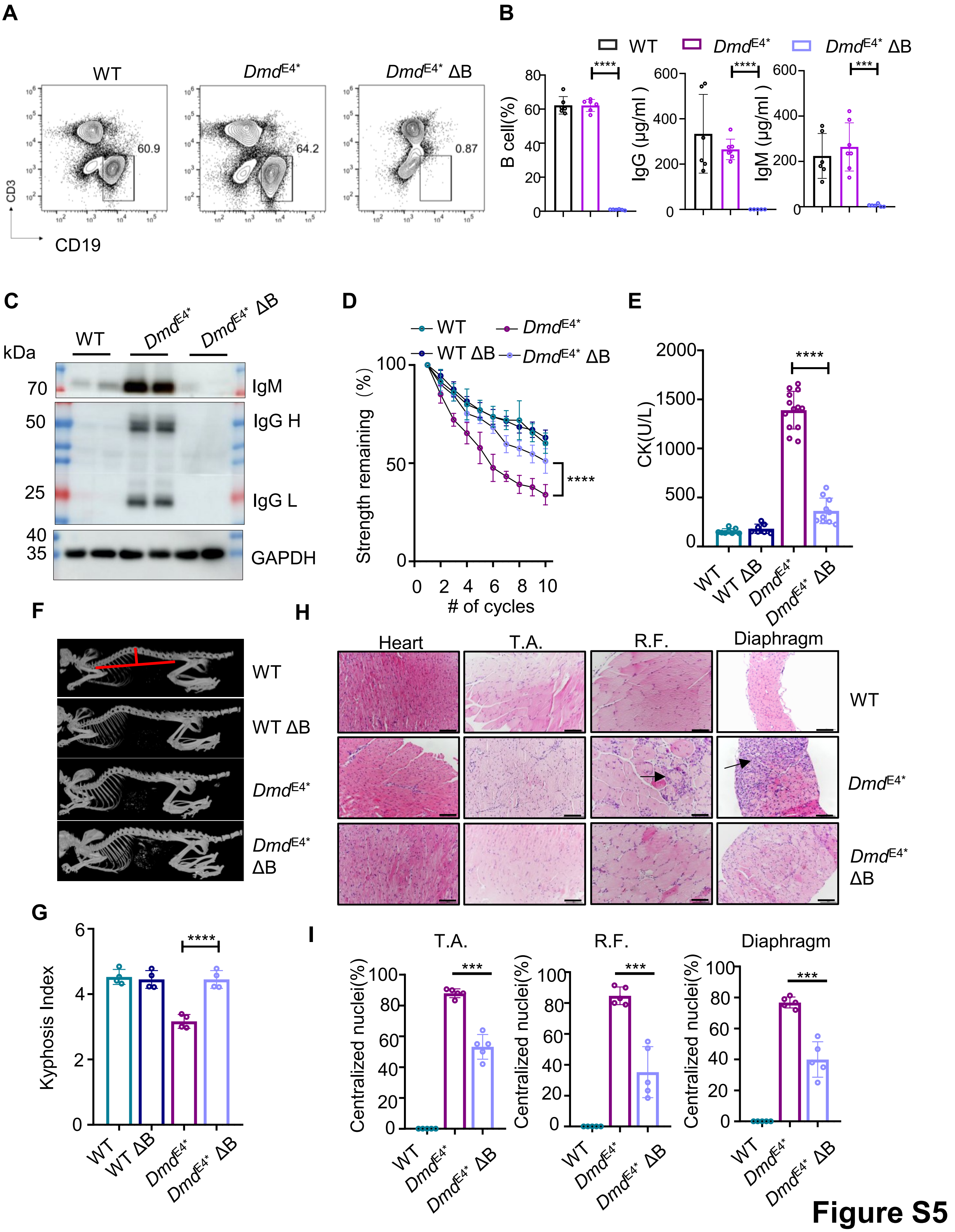

A
WT
DmdE4*
DmdE4* ΔB
60.9
64.2
0.87
CD3
CD19
B
WT
DmdE4*
DmdE4* ΔB
B cell(%)
IgG (µg/ml)
IgM (µg/ml)
****
****
***
C
kDa
WT
DmdE4*
DmdE4* ΔB
70
50
25
40
35
IgM
IgG H
IgG L
GAPDH
D
WT
DmdE4*
WT ΔB
DmdE4* ΔB
Strength remaining (%)
### of cycles
****
E
CK(U/L)
****
WT
WT ΔB
DmdE4*
DmdE4* ΔB
F
WT
WT ΔB
DmdE4*
DmdE4* ΔB
H
Heart
T.A.
R.F.
Diaphragm
WT
DmdE4*
DmdE4* ΔB
G
Kyphosis Index
****
WT
WT ΔB
DmdE4*
DmdE4* ΔB
I
T.A.
R.F.
Diaphragm
Centralized nuclei(%)
***
WT
DmdE4*
DmdE4* ΔB


**Figure S5**

**Fig. S5. Improvement of muscular dystrophy in *Dmd*$^{E4*}$ ∆B mice**

(**A**). Loss of B cells in the *Dmd*$^{E4*}$ ∆B mice. Percentages of B cells ($CD19^+CD45^+$) in PBMC were determined using flow cytometry. Data are either representative (left) or summary (right) of six mice in each group.

(**B**). Loss of B cells, IgG and IgM in the *Dmd*$^{E4*}$ ∆B mice. Serum IgG and IgM levels were determined using enzyme-linked immunosorbent assay (ELISA). Data are summary of six mice in each group.

(**C**). Lack of antibody accumulation in the skeletal muscles of *Dmd*$^{E4*}$ ∆B mice. Tissue lysate from T.A. muscles of WT, *Dmd*$^{E4*}$ and *Dmd*$^{E4*}$ ∆B mice were analyzed by immunoblotting for IgM and IgG. Data are representative of five mice in each group.

(**D**). Enhanced skeletal muscle functions in 2-month-old *Dmd*$^{E4*}$ ∆B mice. 10 whole-body grip strength tests were conducted with a short interval (10 seconds) between each test, and the reduction in strength was calculated by normalized to the maximal grip strength. Data are summarized from five mice in each group.

(**E**). Serum CK levels in 2-month-old WT, WT ∆B, *Dmd*$^{E4*}$, and *Dmd*$^{E4*}$ ∆B mice were determined. Data are summary of at least seven mice in each group. Each dot represents one individual mouse.

(**F**). Representative micro-CT images showing spinal curvature of 8-week-old WT, WT ∆B, *Dmd*$^{E4*}$, and *Dmd*$^{E4*}$ ∆B mice. Data are representative of four mice in each group.

(**G**). As illustrated in (**F**), Kyphosis indexes (KI) were calculated as reported (*34*). Each dot represents one individual mouse (n=4 in each group).

(**H**) Hematoxylin and eosin (H&E) staining of various muscle tissues from *Dmd*$^{E4*}$ ∆B and *Dmd*$^{E4*}$ mice at 8 weeks of age. T.A.: tibialis anterior, R.F.: rectus femoris. Arrows indicate areas with intensive immune cell infiltration. Scale bars: 100 μm. Data are representative of five mice in each group.

(**I**). Percentages of myofibers with centralized nuclei were calculated from 8-week-old WT, *Dmd*$^{E4*}$ and *Dmd*$^{E4*}$∆B mice. Data are a summary of five mice in each group.

Error bars stand for the standard deviation of the mean. *** $p<0.001$, **** $p<0.0001$ in two-tailed Student's *t* test (**B, E**, **G, I**) or two-way ANOVA test (**D**).

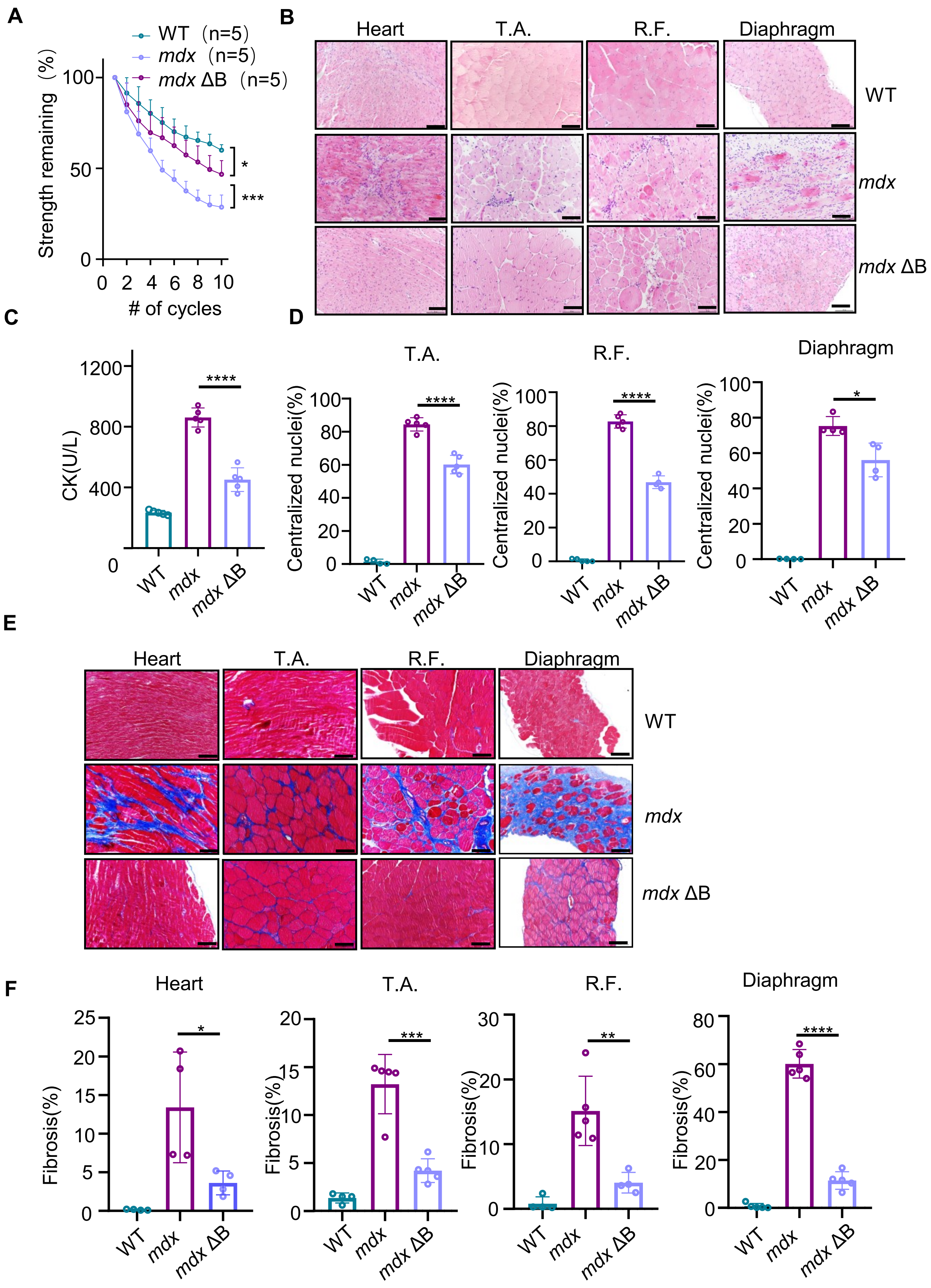
A
WT (n=5)
mdx (n=5)
mdx ΔB (n=5)
Strength remaining (%)
### of cycles
B
Heart
T.A.
R.F.
Diaphragm
WT
mdx
mdx ΔB
C
CK(U/L)
D
Centralized nuclei(%)
E
F
Fibrosis(%)

**Figure S6**

**Fig. S6. Improvement of muscular dystrophy in *mdx* ΔB mice**

(**A**). Improved skeletal muscle functions in 2-month-old *mdx* ΔB mice. 10 whole-body grip strength tests were conducted with a short interval (10 seconds) between each test, and the reduction in strength was calculated by normalized to the maximal grip strength. Data are summarized from 5 mice in each group. **** $p<0.0001$ in two-way comparison ANOVA test.

(**B**). Hematoxylin and eosin (H&E) staining of various muscle tissues from *mdx* ΔB and *mdx* mice at 12 months of age. T.A.: tibialis anterior, R.F.: rectus femoris. Data are representative of four to five mice in each group. Scale bars: 100 μm.

(**C**). Serum CK levels in 12-month-old *mdx* ΔB mice were compared to those in age-matched WT and *mdx* mice. Data are summarized from 5 mice in each group.

(**D**). Percentages of myofibers with centralized nuclei were determined in different skeletal muscles of 12-month-old WT, untreated *mdx* and *mdx* ΔB mice. Data are a summary from four or five mice in each group. Each dot represents one individual mouse.

(**E,F**). Masson's staining of various muscle tissues from *mdx* ΔB, *mdx* and littermate control mice (WT) at 12-month-old -weeks of age. Data are representatives (**E**) or summary (**F**) of four to five mice in each group. Each dot represents one individual mouse. Scale bars: 100 μm.

Error bars stand for the standard deviation of the mean. * $p<0.05$, ** $p<0.01$, *** $p<0.001$, **** $p<0.0001$ in two-tailed Student's *t* test (**C**, **D, F**) or multiple comparison ANOVA test (**A**).

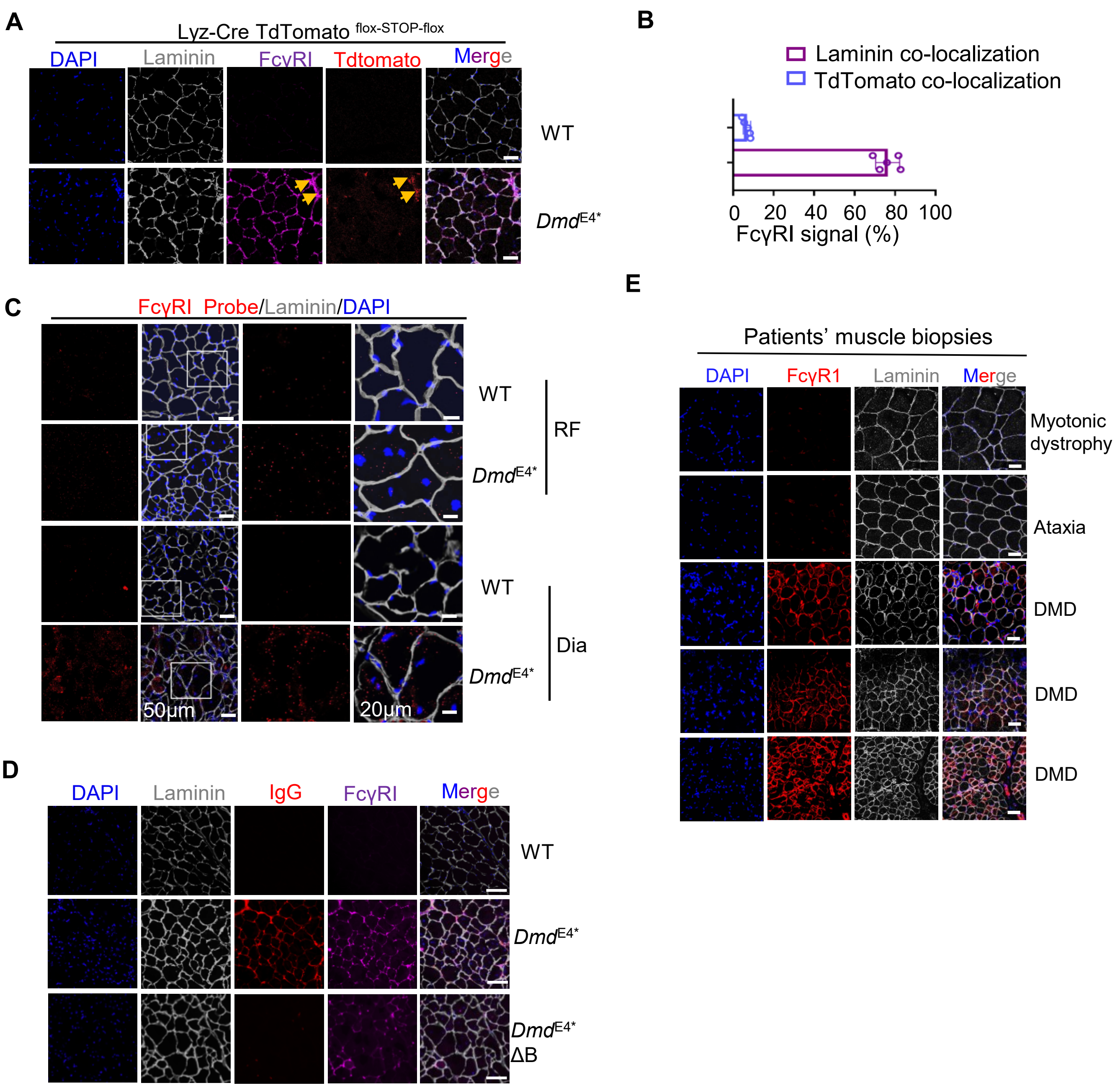

A
Lyz-Cre TdTomato flox-STOP-flox
DAPI
Laminin
FcγRI
Tdtomato
Merge
WT
DmdE4*
B
Laminin co-localization
TdTomato co-localization
0
20
40
60
80
100
FcγRI signal (%)
C
FcγRI Probe/Laminin/DAPI
WT
DmdE4*
RF
WT
DmdE4*
Dia
50μm
20μm
D
DAPI
Laminin
IgG
FcγRI
Merge
WT
DmdE4*
DmdE4* ΔB
E
Patients' muscle biopsies
DAPI
FcγR1
Laminin
Merge
Myotonic dystrophy
Ataxia
DMD
DMD
DMD


**Figure S7**

**Fig. S7. Ectopic presence of FcγR1 on myofibers recruits IgG to dystrophic myofibers.**

(**A,B**). T.A. muscle from Lyz-Cre TdTomato $^{LSL}$ *Dmd*$^{E4*}$ mice was analyzed for levels of FcγR1 and TdTommato by immunofluorescence staining. WT Lyz-Cre TdTomato $^{LSL}$ mice were included as controls. Arrows indicate Tdtommato$^{+}$ cells, which also showed strong FcγR1 expression. Data are representative (**A**) or summary (**B**) of five mice in each group. Note, most FcγR1 signaling was derived from laminin-expressing cells, rather than TdTomato+ cells. Scale bars represent 50 μm.

**(C).** Representative RNA scope analysis of FcγR1 mRNA in R.F and Diaphram. Data are representative of five mice in each group. Scale bars represent 50 μm (left panels) and 20 μm (right panels) respectively.

(**D**). Expression of FcγR1 was sustained in the myofibers of *Dmd*$^{E4*}$ ΔB mice, albeit with a minor reduction. T.A. muscle from eight-week-old *Dmd*$^{E4*}$ΔB mice was analyzed for FcγR1 with immunofluorescence staining. Age matched WT mice and *Dmd*$^{E4*}$ were included as controls. Data are representative or summary of four mice in each group.

(**E**). Elevated FcγR1 expression on myofibers from patients with DMD. Muscle biopsies from patients with DMD were examined for the expression of FcγR1 with immunofluorescence staining. Scale bars represent 20 μm.

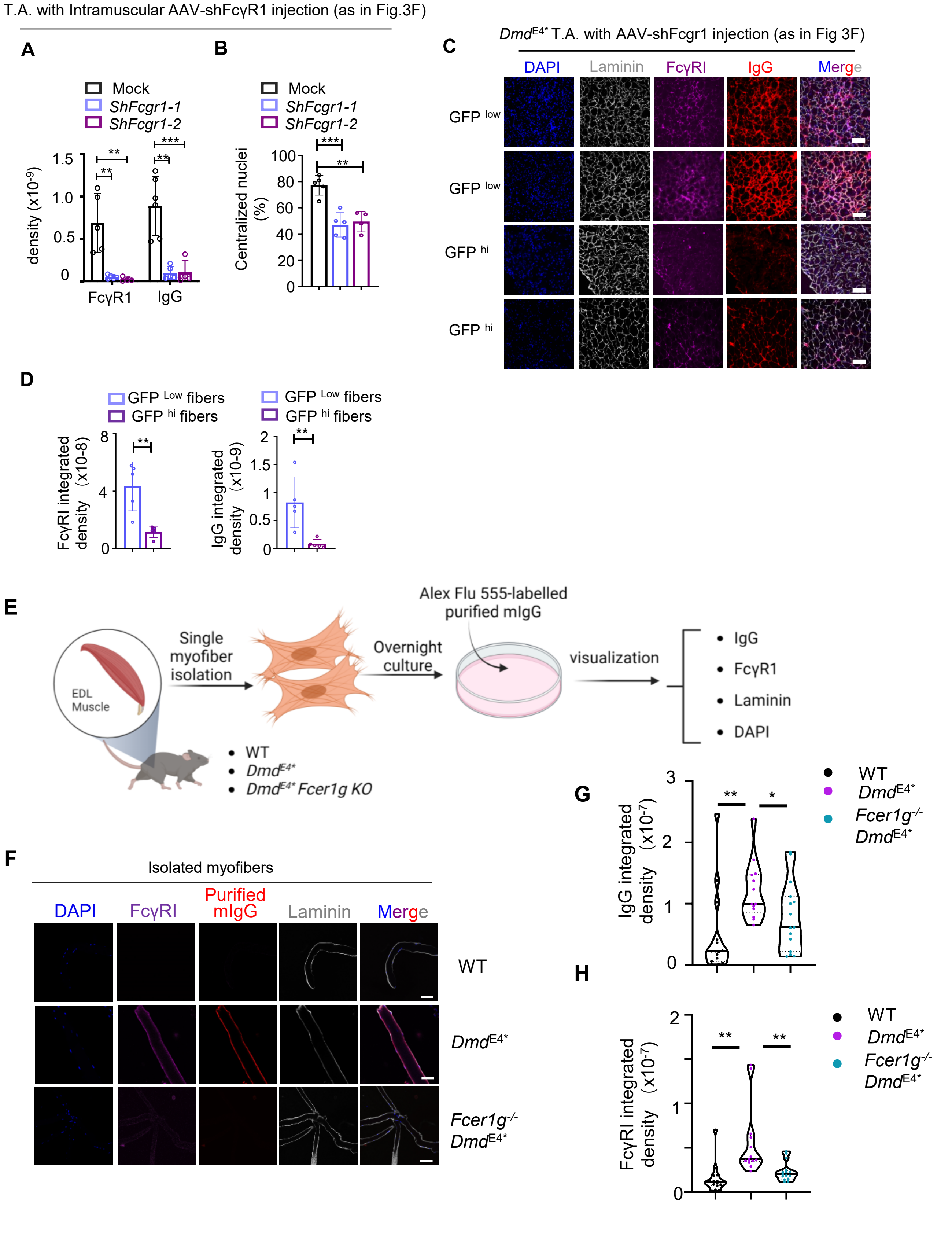
T.A. with Intramuscular AAV-shFcγR1 injection (as in Fig.3F)
A
B
Mock
ShFcgr1-1
ShFcgr1-2
density (x10-9)
FcγR1
IgG
Centralized nuclei (%)
C
DmdE4* T.A. with AAV-shFcgr1 injection (as in Fig 3F)
DAPI
Laminin
FcγRI
IgG
Merge
GFP low
GFP low
GFP hi
GFP hi
D
GFP Low fibers
GFP hi fibers
FcγRI integrated density (x10-8)
IgG integrated density (x10-9)
E
EDL Muscle
Single myofiber isolation
Overnight culture
Alex Flu 555-labelled purified mIgG
visualization
IgG
FcγR1
Laminin
DAPI
WT
DmdE4*
DmdE4* Fcer1g KO
F
Isolated myofibers
DAPI
FcγRI
Purified mIgG
Laminin
Merge
WT
DmdE4*
Fcer1g-/- DmdE4*
G
IgG integrated density (x10-7)
WT
DmdE4*
Fcer1g-/- DmdE4*
H
FcγRI integrated density (x10-7)

**Figure S8**

**Fig. S8. Ectopic presence of FcγR1 on myofibers recruits IgG to dystrophic myofibers.**

(**A-B**). As in **Figure 3G**, levels of FcγR1 and IgG accumulation of myofibers were determined using Image J (**A**). Myofiber damage was determined by identifying myofibers with centralized nuclei (**B**). Scale bars represent 100 μm. Data are summary of four to six mice in each group, and each dot represents one individual mouse.

(**C,D**). As in **Figure 3F**, the T.A. muscle injected with AAV containing sh-Fcgr1 and GFP was analyzed by immunofluorescence staining. GFP $^{hi}$ and GFP $^{low}$ myofibers were identified, and levels of IgG accumulation and FcγR1 were analyzed(**D**). High-magnification view of the GFP $^{hi}$ and GFP $^{low}$ myofibers(**C**). As in (**A**), quantitative analysis of IgG and FcγRI staining intensities in the top 20% (GFP$^{hi}$) and bottom 20% (GFP$^{low}$) of myofibers.

(**E-H**). Myofibers were isolated from EDL(Extensor digitorum longus) of indicated mice and then cultured overnight. Subsequently, the cells were incubated with APC-labeled anti-FcγR1, Alexa Fluor 555-labeled-mIgG, and Alexa Fluor 488-labeled-anti-Laminin on ice overnight(**E**). The fluorescence intensity of each myofiber was determined. Data are either representative (**F**) or summary (**G, H**) of three independent experiments, and each dot represents one myofiber. Scale bars represent 100 μm.

Error bars stand for the standard deviation of the mean. * p<0.05, ** p<0.01, *** p<0.001 in two-tailed Student's *t* test (**A**, **B, D, G, H**).

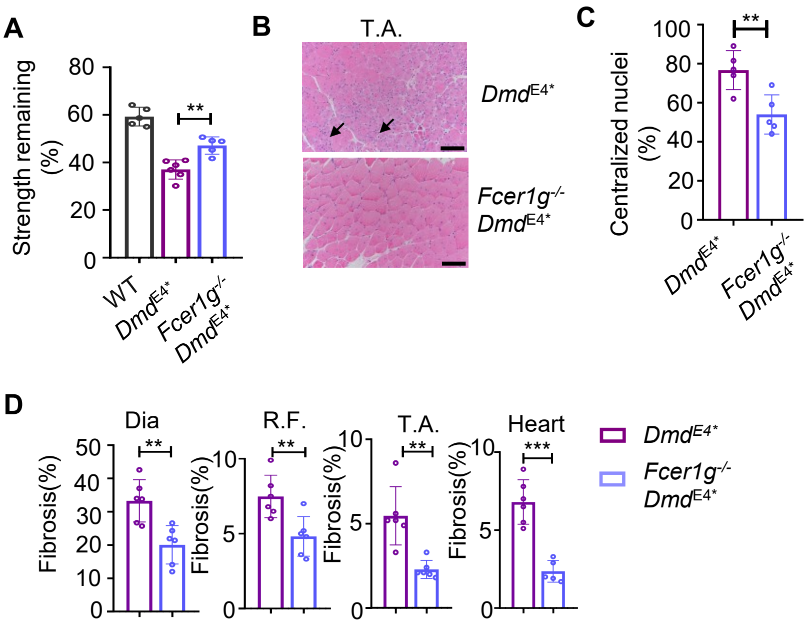
A
Strength remaining (%)
80
60
40
20
0
**
WT
DmdE4*
Fcer1g-/- DmdE4*
B
T.A.
DmdE4*
Fcer1g-/- DmdE4*
C
Centralized nuclei (%)
100
80
60
40
20
0
**
DmdE4*
Fcer1g-/- DmdE4*
D
Dia
R.F.
T.A.
Heart
Fibrosis(%)
50
40
30
20
10
0
**
10
5
0
**
10
5
0
**
10
5
0
***
DmdE4*
Fcer1g-/- DmdE4*

**Figure S9**

**Fig. S9. Genetic ablation of FcγR1 mitigates muscular dystrophy in *Dmd*$^{E4*}$ mice**

(**A**). Ablation of FcγR1 alleviates muscular dystrophy. 6-week-old *Fcer1g*$^{-/-}$ *Dmd*$^{E4*}$ mice and *Dmd*$^{E4*}$ mice were analyzed for remaining grip strength was determined after ten repeated tests. Data are a summary of five mice in each group, and each dot represents one individual mouse.

(**B**). Ablation of FcγR1 diminished tissue inflammation in *Dmd*$^{E4*}$ mice. T.A. muscles from six to seven-week-old *Fcer1g*$^{-/-}$ *Dmd*$^{E4*}$ mice and age-matched *Dmd*$^{E4*}$ mice were analyzed by H&E staining. Arrows indicate areas with inflammatory cell infiltration. Data are representative of 5 mice per group. Scale bars represent 100 μm.

(**C**). Myofibers with centralized nuclei were determined in T.A. muscles. Data are summary of 5 mice per group.

(**D**). As in **Figure 3J,** seven-month-old *Fcer1g*$^{-/-}$ *Dmd*$^{E4*}$ mice and *Dmd*$^{E4*}$ mice were analyzed for fibrosis of various muscle tissues with Masson's staining. Data are a summary of at least five mice in each group, and each dot represents one individual mouse.

Error bars stand for the standard deviation of the mean. * $p<0.05$, ** $p<0.01$, *** $p<0.001$ in two-tailed Student's *t* test (**A**, **C, D**).

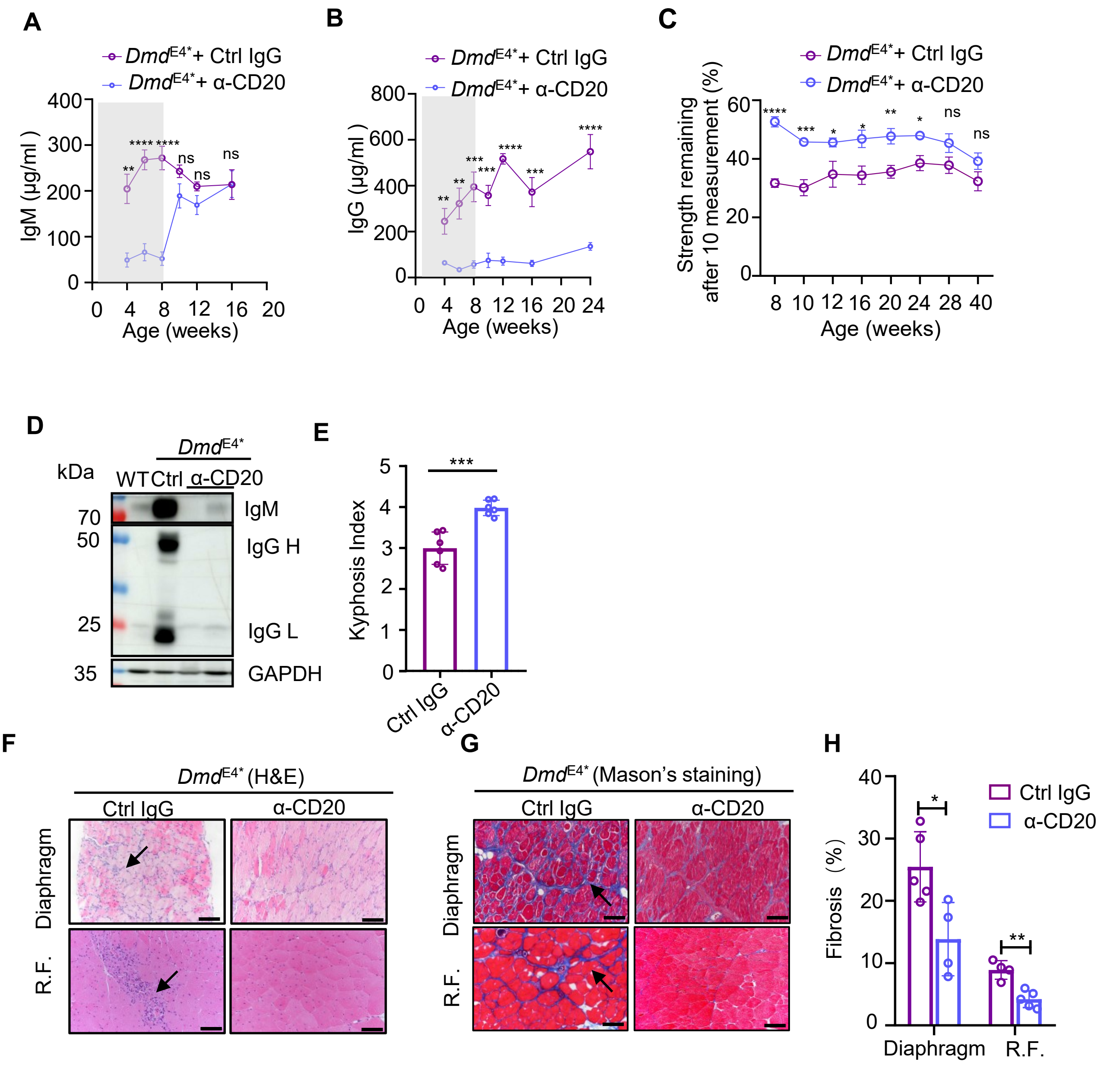
A
DmdE4*+ Ctrl IgG
DmdE4*+ α-CD20
IgM (μg/ml )
Age (weeks)
B
IgG (μg/ml )
C
Strength remaining after 10 measurement (%)
D
kDa
WT Ctrl α-CD20
IgM
IgG H
IgG L
GAPDH
E
Kyphosis Index
Ctrl IgG
α-CD20
F
DmdE4* (H&E)
Diaphragm
R.F.
G
DmdE4* (Mason's staining)
H
Fibrosis (%)

**Figure S10**

**Fig. S10. Depleting B cells using anti-CD20 antibody alleviates muscular dystrophy in DMD mice.**

(**A**) Anti-CD20 treatment in neonatal *Dmd*$^{E4*}$ mice led to transient depletion of immunoglobulin M (IgM). One-week old *Dmd*$^{E4*}$ mice were treated with anti-CD20 or control IgG as depicted in **Figure 5A**. Serum IgM levels were determined using enzyme-linked immunosorbent assay (ELISA). The shaded area indicates the period during which the anti-CD20 treatment was administered. Data are summarized from five to six mice in each group.

(**B**). Anti-CD20 treatment in neonatal *Dmd*$^{E4*}$ mice led to hypogammaglobulinemia. As in (**A**), serum IgG levels were determined using ELISA. Data are summarized from five to six mice in each group.

(**C**) Sustained improvement of skeletal muscle functions following neonatal anti-CD20 treatment. To assess muscle strength, 10 grip tests of combined forelimb and hindlimb strength were conducted every 2-4 weeks, with a short interval (10 seconds) between each test. The remaining strength in the last cycle was calculated by normalizing to the maximal grip strength. Data are summarized from five mice in each group.

(**D**). Eight weeks following the anti-CD20 treatment, levels of IgG and IgM in the T.A. muscle were analyzed using immunoblotting. Data are summarized from three to six mice in each group.

(**E**). Anti-CD20 treatment rescued the *Dmd*$^{E4*}$ mice from kyphosis. Kyphosis indexes (KI) were determined using μCT. Data are a summary of six mice in each group.

(**F**). H&E staining of R.F. and diaphragm from *Dmd*$^{E4*}$ mice treated with anti-CD20 or control mouse IgG. Arrows indicate the area with intense inflammatory cell infiltration. Data are representative of four mice in each group. Scale bars represent 100 μm.

(**G**). Fibrosis was determined using Masson's staining of the diaphragm and R.F. from *Dmd*$^{E4*}$ mice treated with anti-CD20 or control mouse IgG as shown in (**A**). Data are representative of four to five mice in each group. Scale bars represent 100 μm.

(**H**). As in (**G**), fibrotic tissues were quantified based on Masson's staining in the R.F. and Diaphragm. Data are summarized from four or five mice in each group.

Error bars stand for the standard deviation of the mean. * p<0.05, ** p<0.01, *** p<0.001, **** p<0.0001 in two-tailed Student's *t* test (**A**, **B**, **C, E, H**).

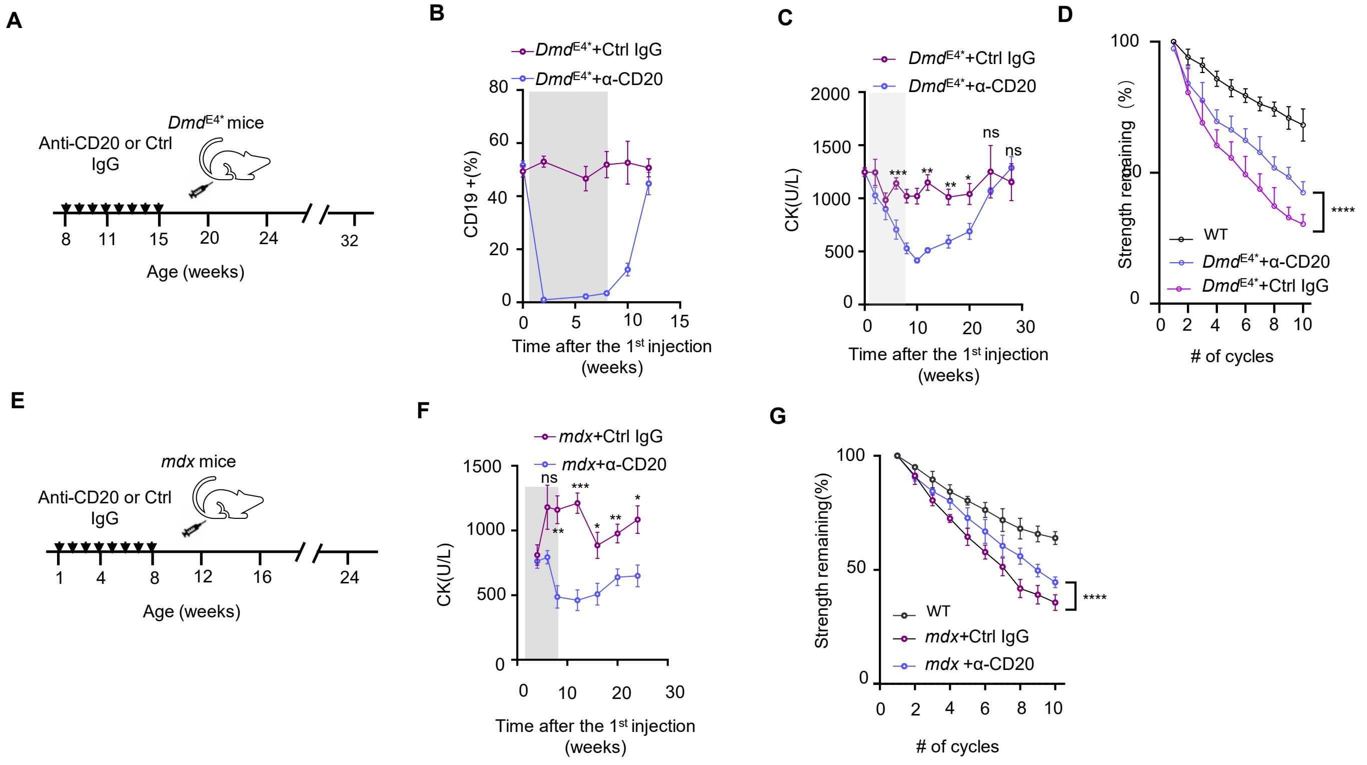
A
Anti-CD20 or Ctrl IgG
DmdE4* mice
Age (weeks)
B
DmdE4*+Ctrl IgG
DmdE4*+α-CD20
CD19 +(%)
Time after the 1st injection (weeks)
C
DmdE4*+Ctrl IgG
DmdE4*+α-CD20
CK(U/L)
Time after the 1st injection (weeks)
D
Strength remaining (%)
WT
DmdE4*+α-CD20
DmdE4*+Ctrl IgG
### of cycles
E
mdx mice
Anti-CD20 or Ctrl IgG
Age (weeks)
F
mdx+Ctrl IgG
mdx+α-CD20
CK(U/L)
Time after the 1st injection (weeks)
G
Strength remaining(%)
WT
mdx+Ctrl IgG
mdx +α-CD20
### of cycles

**Figure S11**

**Fig. S11. Anti-CD20 treatment alleviates muscular dystrophy in adult *Dmd*$^{E4*}$ and *mdx* mice.**

(**A**). Schematic of using anti-CD20 antibody to deplete B cells in 8-week-old *Dmd*$^{E4*}$ mice. 8-week-old *Dmd*$^{E4*}$ mice were injected with anti-mouse CD20 (2 mg/kg) or control mouse IgG once a week for eight weeks.

(**B**). Levels of circulating B cells in PBMC were determined by flow cytometry. Data are summarized from 5 to 6 mice in each group.

(**C**). Serum CK levels were determined prior to the treatment (week 0), and every 2-4 weeks thereafter. Gray areas indicate the time that the mice received the antibody treatment. Each dot represents one individual mouse (five to six mice in each experiment). Note that CK level reduction was sustained for at least 3 months after the last anti-CD20 treatment.

(**D**). As in **Figure 5D**, anti-CD20 treatment prevented contraction-induced fatigue in adult *Dmd*$^{E4*}$ mice. Two weeks after the last injection, 10 grip tests of combined forelimb and hindlimb strength were conducted with a short interval (10 seconds) between each test. The remaining strength in the last cycle was calculated by normalizing to the maximal grip strength. Data are summarized from 5 mice in each group. Each dot represents one individual mouse.

(**E-G**). Anti-CD20 treatment alleviates muscular dystrophy in neonatal *mdx* mice.

(**E**). Schematic of using anti-CD20 antibody to deplete B cells in 1-week-old *mdx* mice. 1-week-old *mdx* mice were injected with anti-mouse CD20 (2 mg/kg) or control mouse IgG once a week for eight weeks.

(**F**). Levels of serum CK were determined in the indicated time. Gray area shows the time window that the mice received the antibody. Data are summarized from five mice in each group. Each dot represents one individual mouse.

(**G**). Eight weeks after the last treatment, 10 grip tests of combined forelimb and hindlimb strength were conducted with a short interval (10 seconds) between each test. The remaining strength in the last cycle was calculated by normalizing to the maximal grip strength. Data are summarized from five mice in each group.

Error bars stand for the standard deviation of the mean. * p<0.05, ** p<0.01, *** p<0.001, **** p<0.0001, ns, not statistically significant in two-tailed Student's *t* test (**B, C, F**) or two-way ANOVA test (**D, G**).

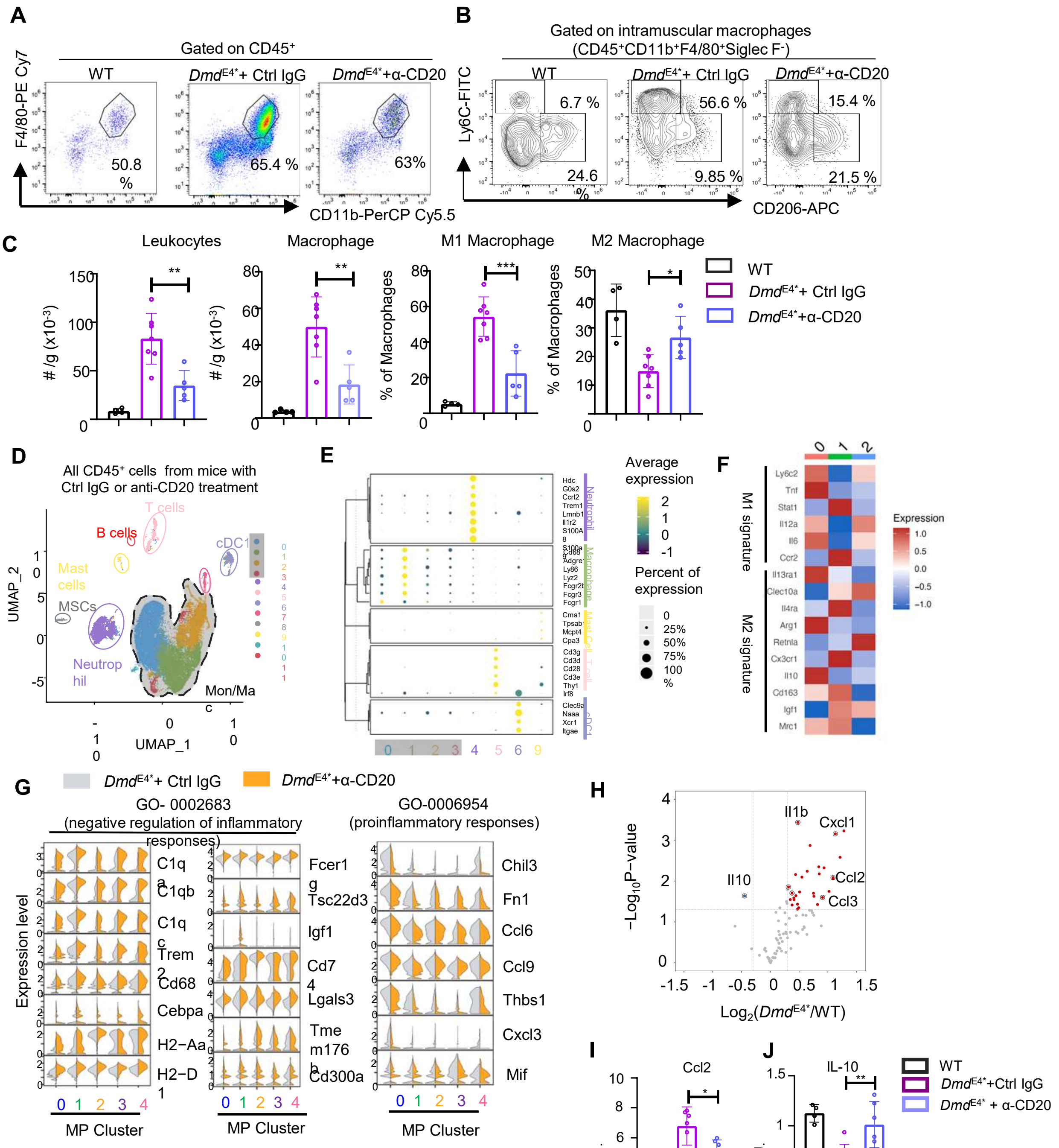

A
Gated on CD45+
WT
DmdE4*+ Ctrl IgG
DmdE4*+α-CD20
F4/80-PE Cy7
CD11b-PerCP Cy5.5
50.8 %
65.4 %
63%
B
Gated on intramuscular macrophages
(CD45+CD11b+F4/80+Siglec F-)
Ly6C-FITC
CD206-APC
6.7 %
24.6 %
56.6 %
9.85 %
15.4 %
21.5 %
C
Leukocytes
Macrophage
M1 Macrophage
M2 Macrophage
# /g (x10-3)
% of Macrophages
WT
DmdE4*+ Ctrl IgG
DmdE4*+α-CD20
D
All CD45+ cells from mice with Ctrl IgG or anti-CD20 treatment
B cells
T cells
cDC1
Mast cells
MSCs
Neutrophil
Mon/Mac
UMAP_1
UMAP_2
E
Average expression
Percent of expression
F
M1 signature
M2 signature
Expression
G
DmdE4*+ Ctrl IgG
DmdE4*+α-CD20
GO- 0002683 (negative regulation of inflammatory responses)
GO-0006954 (proinflammatory responses)
Expression level
C1qa
C1qb
C1qc
Trem2
Cd68
Cebpa
H2−Aa
H2−D1
Fcer1g
Tsc22d3
Igf1
Cd74
Lgals3
Tmem176b
Cd300a
Chil3
Fn1
Ccl6
Ccl9
Thbs1
Cxcl3
Mif
MP Cluster
H
Il1b
Cxcl1
Il10
Ccl2
Ccl3
−Log10P−value
Log2(DmdE4*/WT)
I
Ccl2
A.U.
J
IL-10
WT
DmdE4*+Ctrl IgG
DmdE4* + α-CD20
